# *M. tuberculosis* meets European Lead Factory – identification and structural characterization of novel Rv0183 inhibitors using X-ray crystallography

**DOI:** 10.1101/2025.06.12.659322

**Authors:** Lina Riegler-Berket, Laura Gödl, Nakia Polidori, Philipp Aschauer, Christoph Grininger, Gareth Prosser, Jörg Lichtenegger, Theo Sagmeister, Lena Parigger, Christian Gruber, Norbert Reiling, Monika Oberer

## Abstract

Tuberculosis, caused by *Mycobacterium tuberculosis* (Mtb), remains a leading cause of mortality worldwide. Proteins involved in lipid metabolism, such as the monoacylglycerol lipase Rv0183, play critical roles during both the active and dormant phases of Mtb and present novel targets for therapeutic intervention. Through high-throughput screening at the European Lead Factory, we identified a novel chemotype characterized by a hydroxypyrrolidine ring, which demonstrated potent inhibition of Rv0183 and promising results in whole cell bacterial studies. Subsequent co-crystallization studies of this chemotype with Rv0183 revealed non-covalent interactions within the lipase’s binding pocket, elucidating the inhibitory mechanism. Comparative analysis, augmented by AI-driven 3D-point-cloud approaches, distinguished Rv0183’s ligand-binding cavity from that of human monoacylglycerol lipase, implying the possibility for species-selective inhibition. This selectivity was further supported by molecular docking simulations which validated the experimental binding affinities and predicted strong, specific binding modes. Our study presents not only the structural basis for the inhibition of Rv0183 by these novel hydroxypyrrolidine-based inhibitors but also demonstrates the utility of integrating computational and empirical methods to achieve species-specific targeting. This approach could minimize off-target effects in humans, marking a significant step toward developing more effective antitubercular therapies. The potential to selectively inhibit Mtb in its dormant state could lead to treatments that prevent the persistence and resurgence of the disease, addressing a crucial gap in the fight against tuberculosis.

## Introduction

Tuberculosis (TB) is a significant health threat due to its high mortality rate and the challenges associated with its treatment and prevention. In 2023, a total of 1.25 million people died from TB and another 10.8 million people fell ill with TB worldwide. These numbers make *Mycobacterium tuberculosis* (Mtb), the causative agent for TB, again the leading single infectious killer after COVID-19 had replaced it for three years ^1^. Accordingly, Mtb has been added to the updated Bacterial Priority Pathogens List in 2024 ^2^. WHO has called for a 90% reduction in TB-death for 2030 in its ‘The End TB Strategy’ ^3^. The disease’s ability to go in latency, coupled with the risk of inadequate treatment compliance leads to more and more drug resistant strains. Most antibiotics used to treat tuberculosis work on replicating bacteria ^4,5^. However, a major chance - and challenge - to eradicate TB is the bacterium’s ability to transition from a replicating to a non-replicating state, the latter having less susceptibility to the action of antibiotics and resulting in poor treatment outcomes ^6,7^. To address this issue, it is crucial to identify and target proteins that are active during dormancy ^8,9^. Proteins in lipid metabolism of Mtb have been identified as having great potential as drug targets, due to their important roles in the generation of energy, maintenance of cell integrity, and the synthesis of the cell wall during dormancy ^10–14^.

The Mtb genome encodes a remarkable number of serine hydrolases compared to other bacterial pathogens and humans: they represent 4.4% of the Mtb proteome, whereas this enzyme class represents 1.5-2.2% in other pathogens ^15^ and 1.2% in humans ^16^. Studies from two independent laboratories indicate that the serine hydrolase monoacylglycerol lipase Rv0183 remains active during the dormant/persistent and the reactivation stage ^17–19^.

Rv0183 was found in the cell wall and in the culture medium as secreted lipase ^20,21^. Sera of TB patients show a humoral immune response with IgG antibodies specific for Rv0183 ^11^. The lipase has been associated with the degradation of lipid droplets present in adipocytes or foamy macrophages, resulting in the generation of free fatty acids that may be used as energy supply during dormancy ^14,20,22,23^. Rv0183 seems to be important but not essential, as clarified by high density mutagenesis studies of Sassetti et al. in 2003 and by CRISPR interference-based functional genomics method of Bossh et al. in 2021. ^24,25^. Knockouts of the homologous gene *MSMEG_0220* in *M. smegmatis* drastically altered colony morphology and changes susceptibility to antibiotics. This observation suggests that the phenotype change could impact bacterial stability within macrophages and alter intercellular interactions ^22^. A study of the role of Rv0183 in mouse alveolar macrophages shows that persistent expression of Rv0183 enhances the immune response by increasing the expression of several cytokines which subsequently can cause tissue damage and host cell death ^26,27^. Based on the observations, that i) Rv0183 is actively expressed during dormancy and the active stage of Mtb, ii) loss of the ortholog gene in *M. smegmatis* results in significant changes of cell morphology and drug susceptibility, and iii) Rv0183 modulates the immune response in alveolar macrophages, we study Rv0183 as target for pharmacological inhibition aiming at potential beneficial therapeutic output of individuals infected with Mtb.

In previous studies, we determined the three-dimensional (3D) structure of Rv0183 using X-ray crystallography ^28,29^. Rv0183 has an α/β hydrolase fold harboring a catalytic triad (S110, D226 and H256) with a cap structure that covers the binding pocket. The cap of Rv0183 adopts a Z-like shape typical for lipases with high specificity for monoacylglycerol (MG) ^30^. Based on the ability of the mycobacterial lipase Rv0183 to effectively hydrolyze MGs of different chain lengths, it also has been termed as monoacylglycerol lipase from *M. tuberculosis* (mtbMGL) ^20,28,29^. Rv0183 also hydrolyzes diacylglycerol, yet to a lesser extent than MG; no activity towards triacylglycerols was observed in different laboratories ^20,29^. Our previously reported crystal structures already gave important insights into the nature of the binding pocket: They reveal binding modes and orientation of the acyl-chain of the MG substrate and give first indications, that selective inhibition of Rv0183 over human MGL is possible upon analysis of similar and distinct features of the binding pocket ^28,29^.

To achieve the goal in identifying inhibitors for Rv0183, we successfully applied at the European Lead Factory (ELF), acting as a shared platform for drug discovery that was funded by the Innovative Medicines Initiative (IMI) and the Innovative Health Initiative. The ELF library houses a high-quality compound library of unique chemical compounds with drug-like properties from pharmaceutical companies, academic partners and medicinal chemists ^31^. There, we pursued a high-throughput screening (HTS) effort to generate effective starting points for our drug discovery efforts against Rv0183. This paper presents the first small molecule inhibitors of Rv0183 that we obtained from this ELF screening campaign. While the complete screening library contained approx. 470 000 compounds, 33 compounds were selected as potential hits and included in the quality hit list (QHL). All compounds of the QHL demonstrated an IC50 in the low µM range/ pIC50 > 5. We co-crystallized three inhibitors named ELF1, ELF5, and ELF8 with Rv0183 and determined high resolution structures. Importantly, these compounds demonstrated growth inhibitory activity against MtbH37Rv when grown with monoolein as a sole carbon source. These three lead compounds share a 3-hydroxypyrrolidin-1yl-methanone moiety which is substituted by different ring systems. Our crystallographic analysis reveals that 3-hydroxypyrrolidin-1-yl-methanone moiety is strategically positioned to engage in a critical interaction in proximity to the catalytic site of Rv0183. Furthermore, we identified interactions of the substituted pyrazole, pyrimidine and oxazole ring systems along the substrate binding pocket. We also employed an innovative 3D-point-cloud based approach to compare the binding cavities of the inhibitor-binding sites in the available structures of Rv0183 and their human orthologs. Our analysis disclosed a discrete and distinctly apart grouping of Rv0183 from the human monoacylglycerol lipase (hMGL). Thus, our study reveals unprecedent insights into structural details of the species-selective inhibition of Rv0183 from *M. tuberculosis*, a potential druggable lipase that is active during dormancy and reactivation of tuberculosis infection.

## Material and Methods

### Materials

*E. coli* strains for protein production and chemicals were purchased from Thermo Fisher Scientific Inc. (Waltham, MA, USA) and Carl Roth GmbH & Co. KG (Karlsruhe, Germany) unless noted otherwise.

### Cloning, expression and purification of Rv0183

Screening at ELF was carried out using wildtype Rv0183 (UniProt Accession code: O07427). All crystallization was carried out with the surface entropy mutant K74A of Rv0183 described in detail before ^29^. The changed residue adopts a surface exposed location after β-strand 3 of the α/β core and is not involved in forming the substrate binding pocket. The variant is fully active in hydrolysing monoacylglycerols, yet displays significantly enhanced crystallization properties^29^. Plasmids of pProExHtb containing the gene encoding Rv0183-K74A for crystallization and Rv0183-wt for activity assays were transformed into BL21 Star™ (DE3) pLysS One Shot™ chemically competent *E. coli* with standard protocols.

A single colony was used for an overnight culture in LB media containing 100 µg/mL ampicillin. 500 mL of LB media containing 100 µg/mL ampicillin were inoculated with 5 mL of overnight culture in a 2 L flask. The culture was grown in a shaking incubator at 37 °C until an OD_600_ greater than 0.7 was reached. Then the Trc promoter was induced with IPTG (final concentration of 1 mM) and incubated overnight at 18°C. The cells were then harvested by centrifugation (40 min, 16000 x g) and pellets containing 1 L fractions were stored at -20°C until purification.

Rv0183-K47A protein was purified according to an established protocol ^29^. 40 mL of buffer A (20 mM Tris-HCl pH 8.0, 150 mM NaCl) was added to a 1 L frozen cell pellet. After thawing on ice, the pellet was homogenized using a dispersing instrument (IKA (Staufen, Germany), T 18 digital ULTRA-TURRAX). Then the homogenate was lysed, using an ultrasonicator (Bandelin (Berlin, Germany), Sonoplus HD 2070), sonicating it for 30 min at 5x 50% with MS72 2mm tip. After centrifugation at 18000 x g for 30 min the supernatant was filtered using a 0.45 µm filter and then loaded onto a gravity flow 5ml His-Trap column (Quiagen (Hilden, Germany)). After a 100 mL wash step with buffer A, the protein of interest was eluted from the gravity flow column using 60 mL of buffer B (20 mM Tris-HCl pH 8.0, 150 mM NaCl, 250 mM imidazole).

The concentrated elution fraction was loaded onto a size-exclusion column (HiLoad 26/600 Superdex 200 pg (Cytiva (Marlborough, USA), 270 mL bed volume, 2 mL/min flow rate) using buffer A. Fractions of the monomeric protein, eluting at about 224 mL, were pooled and concentrated to 10 mg/ml of protein concentration using Centricons (MilliPore (Billerica, USA). 50 µL fractions were flash frozen and stored at -80°C until further use.

### Cloning, expression and purification of human MGL

We generated and used a soluble variant of human MGL (UniProt Accession code: Q99685) as described before ^32^. This variant has three amino-acid exchanges, namely K36A, L169S and L176S leading to increased solubility and crystallizability while retaining MG-hydrolyzing activity. We introduced these mutations to the hMGL gene in the pProEx-vector using the Q5® site-directed mutagenesis kit according to the manufacturer’s protocol. The primers are listed in Table 1. The mutations were verified by Sanger sequencing. The soluble human MGL variant (shMGL) used for the deselection assays in HTS was expressed and purified with the same protocol as described previously for Rv0183.

**Table 1.**
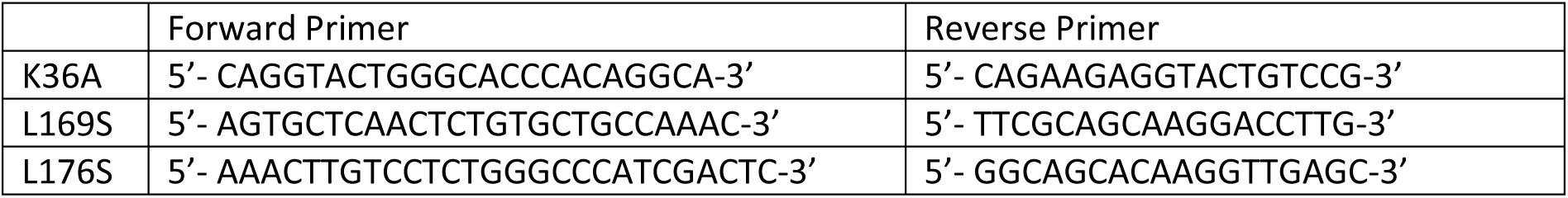
Primers used for generation of the soluble variants of hMGL.

### Screening assays for high throughput screening (HTS)

We conducted an HTS campaign in collaboration with the European Lead Factory at the Pivot Park Screening Center (AB Oss, The Netherlands) and BioAscent Discovery (Newhouse, Lanarkshire, UK) to discover new inhibitors for Rv0183. For the screening campaign, we adopted a previously developed MGL activity assay to the 1536 well-format ^28,33^. The activity assay for the primary screen uses monoolein as a substrate and Free Glycerol Reagent (FGR, Sigma-Aldrich) to read out the glycerol product detected at 540 nm.

The final assay protocol for the primary (P1) and active confirmation assay (C1) was as follows: 0.5 µL of 49.6 nM protein in protein buffer (50 mM Tris-HCl pH 7.4, 150 mM NaCl) was dispensed into Corning 1356 well microplate wells. A volume of 20 nL of compound (or DMSO) was transferred using ECHO acoustic liquid handler to reach a final concentration of 10 µM (final DMSO concentration 0.25 %) and incubated for 30 minutes at room temperature (RT). From the monoolein substrate stock (4 mM monoolein in 0.55 mM potassium phosphate buffer pH 8.0, 5 mM CHAPS), a dilution in free glycerol reagent solution to 320 µM concentration was prepared and a volume of 7.5 µL of substrate dilution was added. After incubation for 40 minutes at room temperature (RT), the absorbance was measured at 540 nm using PHERAstar. For the active confirmation in dose-response curve (DRC) format, the assay conditions were identical, with the exception that 80 nL of compound were added to result in a final concentration of 20 µM to 22 nM in 7 points. Absorption was measured after incubation with substrate for either 40 min or 180 min (C2 and C3, respectively).

The orthogonal assay (O2) was performed in DRC format. Monoolein was used as the substrate and the non-esterified free fatty acid (NEFA) colorimetric assay kit (FUJIFILM Wako Chemicals Europe GmbH) was used to quantify the free fatty acid product. The final assay protocol is identical to that described for the primary and active confirmation assays and in the NEFA kit user manual, with the following exceptions: A volume of 1.5 µL monoolein (300 µM in 0.1 M potassium phosphate buffer) was added to the plate and incubated for 40 min. For the detection of absorbance at 550 nm, a volume of 4 µL detection solution R1 was incubated for 15 minutes, followed by the addition of 2 µL of detection solution R2 and incubation for 15 min at room temperature.

The deselection assays (D2) were conducted in accordance with the protocol for the active confirmation assay in DRC format (40 min incubation time), with the exception that the final assay concentration of hMGL was 1 µM.

A thermal shift assay (TSA) was successfully developed for the Rv0183 protein using the primary assay buffer of 55 mM potassium phosphate at pH 7.3. The assay showed sufficient fluorescence signal and clear derivative peaks in a final volume of 20 µL containing 1.5 µg/well protein (2.34 µM). The experiment was performed using Thermo Scientific Thermal Shift Dye #4461146, which has an excitation/emission wavelength of 580/623 nm. The assay was performed in the dye kit buffer containing 1 M potassium phosphate at pH 7.0. Test samples were prepared on ice in a qPCR plate and subjected to the following cycling conditions: initial incubation at 4°C for 1 minute, followed by heating at a rate of +0.05°C per second until 99°C was reached, then maintaining the temperature at 99°C for 1 minute. The 413 confirmed compounds were tested in triplicate at a single concentration of 10 µM and a final DMSO concentration of 0.25%. In order to define which compounds, affect protein stability, a selection cut-off of greater than 4 times the standard deviation of the mean delta Tm of all Rv0183+DMSO controls across all three test plates was calculated (0.47°C). The high-likelihood binders were identified as those with delta Tm values exceeding 1.5°C, with no interfering fluorescence

### Resynthesis of ELF1, ELF5, ELF8

After screening at the ELF lead factory, compounds were resynthesized (Supplemental Material S1) in sufficient quality and purity as proven with liquid chromatography (LC) and nuclear magnetic resonance (NMR) (Supplemental Material S2).

### Whole cell bacterial experiments

Mtb strain H37Rv (MtbH37Rv) was used for all whole cell bacterial studies. All work was carried out within a Biosafety Level 3 facility, as required by law. MtbH37Rv was routinely propagated in, and minimum inhibitory concentration (MIC) assays performed in, a synthetic growth medium composed of 1 g/L KH_2_PO_4_, 2.5 g/L Na_2_HPO_4_, 0.5 g/L NH_4_Cl, 0.5 g/L Mg_2_SO_4_.7H_2_O, 0.05 g/L ferric ammonium citrate, 0.5 mg/L CaCl_2_, 0.1 mg/L ZnSO_4_, 5 g/L bovine serum albumin (fatty acid free), 0.85 g/L NaCl, and 0.05 % (v/v) tyloxapol. 300 µM monoolein was added as the sole carbon source (from a 300 mM ethanolic stock). The pH of the final medium was adjusted to 6.8 with HCl. MIC assays were performed in clear, round bottomed 96 well plates, with a final volume of 100 µL and a starting bacterial OD600 of 0.001 (∼5000 cells per well). Compounds were tested at 7 different concentrations, consisting of a top concentration of 100 µM and subsequent 2-fold serial dilutions down to 1.56 µM. Final DMSO concentrations did not exceed 0.1 % v/v. Untreated and 2 µM rifampicin (RIF) treated wells were also included to facilitate data analysis. Plates were left to incubate, statically at 37° C in a humidified container, for 7-12 days prior to addition of resazurin (30 µL of a 0.02 % w/v aqueous stock) to all wells.

Red fluorescence (540 nm excitation, 590 nm emission) was recorded after a further 24 hours, and growth inhibition calculated relative to the untreated (100 % growth) and RIF-treated (0 % growth) controls.

### Crystallization and crystal soaking

Crystallization experiments were performed with an ORYX 8 robot (Douglas Instruments (Berkshire, UK). SWISSCI 3 Lens Midi UVXPO-3LENS crystallization plates (SWISSCI AG (Neuheim, Switzerland)) were used for sitting-drop vapor diffusion experiments. Initial hits were found in a Morpheus 3 screen (Molecular Dimensions (Sheffield, UK)) and optimized containing 1.2% cholic acid derivatives mix (3% w/v CHAPS, 3% w/v CHAPSO, 3% w/v sodium glycocholate hydrate, 3% w/v taurocholic acid sodium salt hydrate), 0.1 M Buffer System 2 pH 7.5 (51.8% MOPS, 48.2% HEPES) and 38% Precipitant Mix 4 (25% v/v MPD, 25% PEG 1000, 25% w/v PEG 3350).

Crystallization was carried out in protein crystallization cabinets (RuMED (Laatzen, Germany)) at 20°C. For the setup, 35 µL of crystallization condition were transferred in the reservoir, 0.25 μL of protein solution (10 mg/mL) were mixed with 0.25 μL reservoir solution and 0.1 µL of seed stock. Seed stocks were prepared from Rv0183-K74A crystals in optimized condition by crushing crystals in 30 µL of crystallization condition. A 1:500 dilution was used for the crystallization setup. Crystals grew within two weeks. Solid compounds were soaked into crystal containing drops 2 hours before harvesting and storing in liquid nitrogen.

### Data collection and processing

Diffraction data was collected at the DESY beamline P11 ^34^ and indexed and integrated with XDS ^35^. Access was granted via BAG-20200791 EC. We used CCP4 Cloud (Collaborative Computational Project No. 4) ^36,37^ for data processing. The datasets were cut at the highest resolution limit. For deciding on the cutoff limit we considered completeness (more than 98%) and data redundancy (multiplicity of at least 1.8). Datasets were scaled and merged with the program aimless and the structures were solved by molecular replacement with the program phaser ^38^ using the published structure of the Rv0183 variant K74A (PDB-ID: 7P0Y ^29^) as a search model. The initial models were iteratively refined using refmac5 (CCP4 Cloud) and then models were built manually in Coot (Crystallographic Object-Oriented Toolkit, version 0.9.8.1) ^39^. At this step, the structures of the compounds were built into the structure of the protein using the ligand builder of Coot. The final refinement was performed using phenix.refine ^40^ to refine occupancies of alternative orientations of the inhibitors and side chains. Detailed data processing and structure refinement statistics are summarized in Table 4.

### Off-target analysis

Active-site cavities, represented as 3D point-clouds, of available Rv0183 structures and the human orthologs, which were identified via databases searches with BLAST ^41^ and FoldSeek ^42^, were created with the Catalophore^TM^ technology ^43,44^, employing the LIGSITE algorithm ^45^ with a cutoff value of 5 Å. A comparison of point-clouds by shape and physico-chemical properties was achieved by superimposing and optimizing the alignment of the cavities, so that the matching score was minimized. The binding sites were clustered by their pairwise complementarity of shape and relevant physico-chemical properties by hierarchical clustering with the Python module SciPy ^46^.

### Molecular docking of ELF1, ELF5 and ELF8 into available Rv0183 structures

Molecular docking was performed with AutoDock VINA ^47^ within the CavitOmiX Platform employing Yasara ^48^. The target structures were prepared by removing ligands from the binding site and subsequently minimizing them in the amber03 force field. The docking box was extended by 7 Å around the active-site cavity and dockings were performed 45 times each using the amber03 ^49^ force field. Amino-acid residues which could influence inhibitor binding (L39, H109, S110, L142, V143, V147, V163, G164, V192, L201, L228, H256 and E257) were allowed to experience flexibility during the docking process.

### Visualization

Visualizations of the data were generated using the Matplotlib ^50^ and Seaborn ^51^ libraries in Python. Structural representations were produced with PyMOL (The PyMOL Molecular Graphics System, Version 3.0 Schrödinger, LLC) and the Innophore‘s CavitOmiX plugin (v. 1.0, 2022, Innophore GmbH).

## Results

### High throughput screening selects the best 13 inhibitors out of a library of 470 000 compounds

The selection process from the compound library at ELF included high-throughput (HTS)-assays, orthogonal screening, dose response experiments, de-screening of Rv0183 vs human MGL and triaging. A total of 469,468 compounds in 388 plates were screened for their capacity to inhibit the hydrolytic activity of Rv0183 against monoolein (Figure 1).

**Figure 1.**
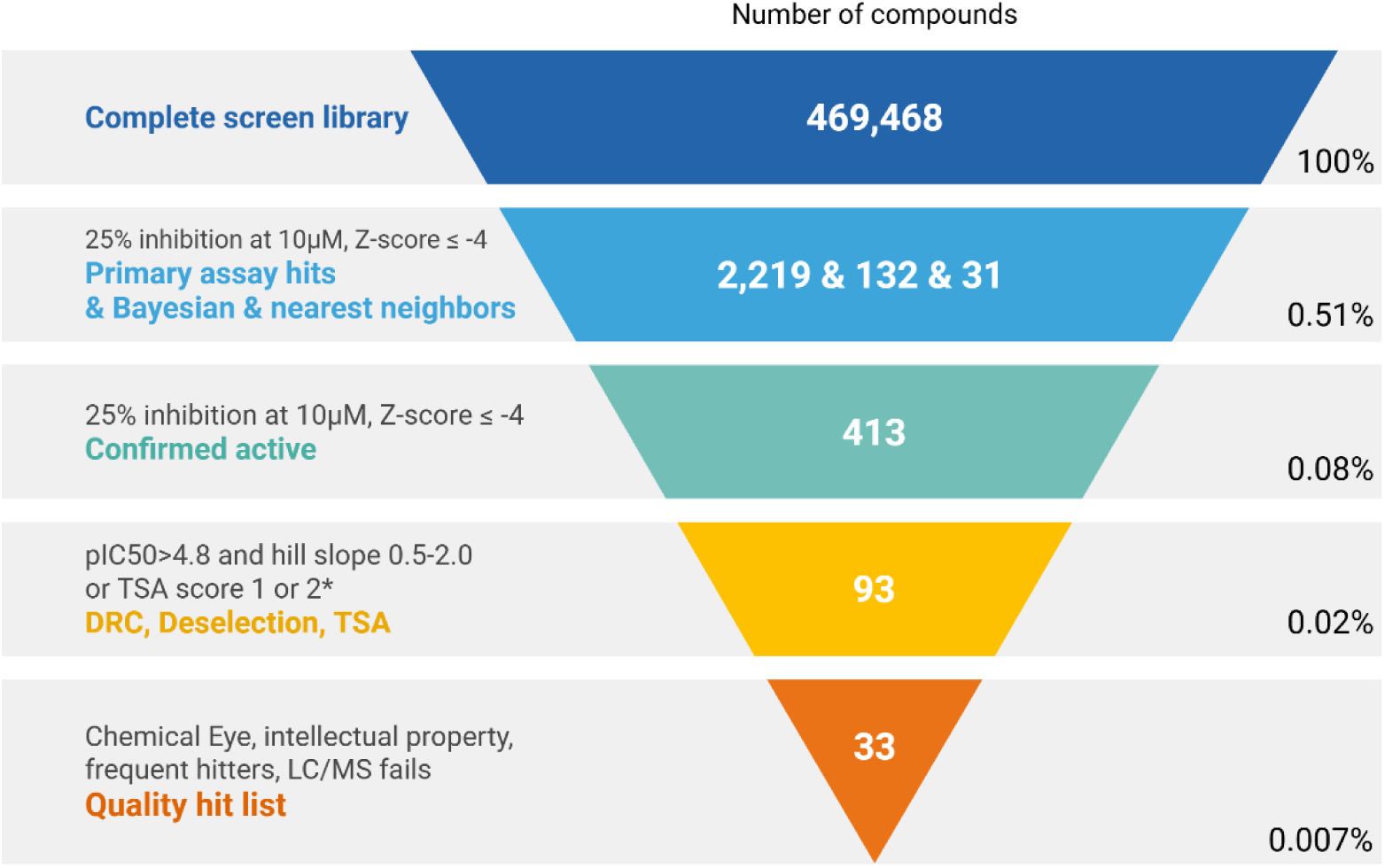
Schematic representation of the selection process in the ELF screening campaign. TSA – Thermal shift assay – Thermal shift either 1.5°C (score 1) or falling between the cut-off (4*SD of the mean Rv0183 + DMSO) and 1.5°C (score 2); DRC – dose response curve.

Screening and validation criteria included a Z-prime greater than 0.6 (actual range: 0.70-0.97) and signal-to-background (S/B) greater than 3 (actual range: 4.9-12.1). This resulted in 2219 compounds that gave a Z-score ≤ -4 and a decreased activity (termed effect) ≥ 25% at an inhibitor concentration of 10 µM. A total of 2382 compounds were selected at this stage (including 132 Bayesians and 31 nearest neighbors) of which 2263 compounds being available for further testing. After retesting of these components in the same assay (Z-prime 0.92-0.94 and S/B range of 10.8-11.3) 413 molecules could be confirmed as active for Rv0183 inhibition with more than 25% inhibition at compound concentrations of 10 µM (Z-score ≤ -4). The selected compounds were tested in dose-response experiments (DRC) using both primary and orthogonal assays. Additionally, the confirmed active compounds were tested in the thermal shift assay (TSA) to provide evidence of target engagement. A total of 93 compounds were selected based on specific criteria, including a pIC50 > 4.8 and a hill slope of 0.5 to 1 in one of the DRC experiments (to exclude compounds with nonspecific effects on the target), or a thermal shift either ≥ 1.5°C (TSA score 1) or falling between the cut-off (4*SD of the mean Rv0183 + DMSO) and 1.5°C (TSA score 2). In addition, a deselection campaign was performed using the human MGL ortholog of Rv0183 at the primary hit list stage. The objective was to select compounds that inhibit Rv0183 with high specificity, while not interfering with hMGL, thus ensuring species-specific activity. However, since no compound showed distinct inhibition activity on hMGL (pIC50 < 4.7) in the primary assay setup, this criterion was not considered further in the selection process. In the next step the list was cleared on the basis of quality control failures (major LC/MS fails), frequent hitters, intellectual property (compounds blocked by the compound owner) and those with unfavorable chemical structures based on structural assessment. Accordingly, 33 compounds were selected for the QHL (Figure 1) including 13 top level, Tier 1, compounds that were recommended for further use, 16 Tier 2 compounds that met some activity criteria but were not directly recommended yet were included to broaden the scope, and 4 Tier 3 compounds that did not meet the activity criteria but were included for structure-activity relationship (SAR) purposes.

Eight of the Tier 1 compounds could be resynthesized, passed the re-testing and quality control, and were subsequently used for follow up crystallization studies. Here we describe the complexed crystal structures of three Tier 1 compounds with Rv0183. The results of the HTS as well as the chemical structures for these compounds are summarized in Figure 1 and Table 2.

**Table 2.**
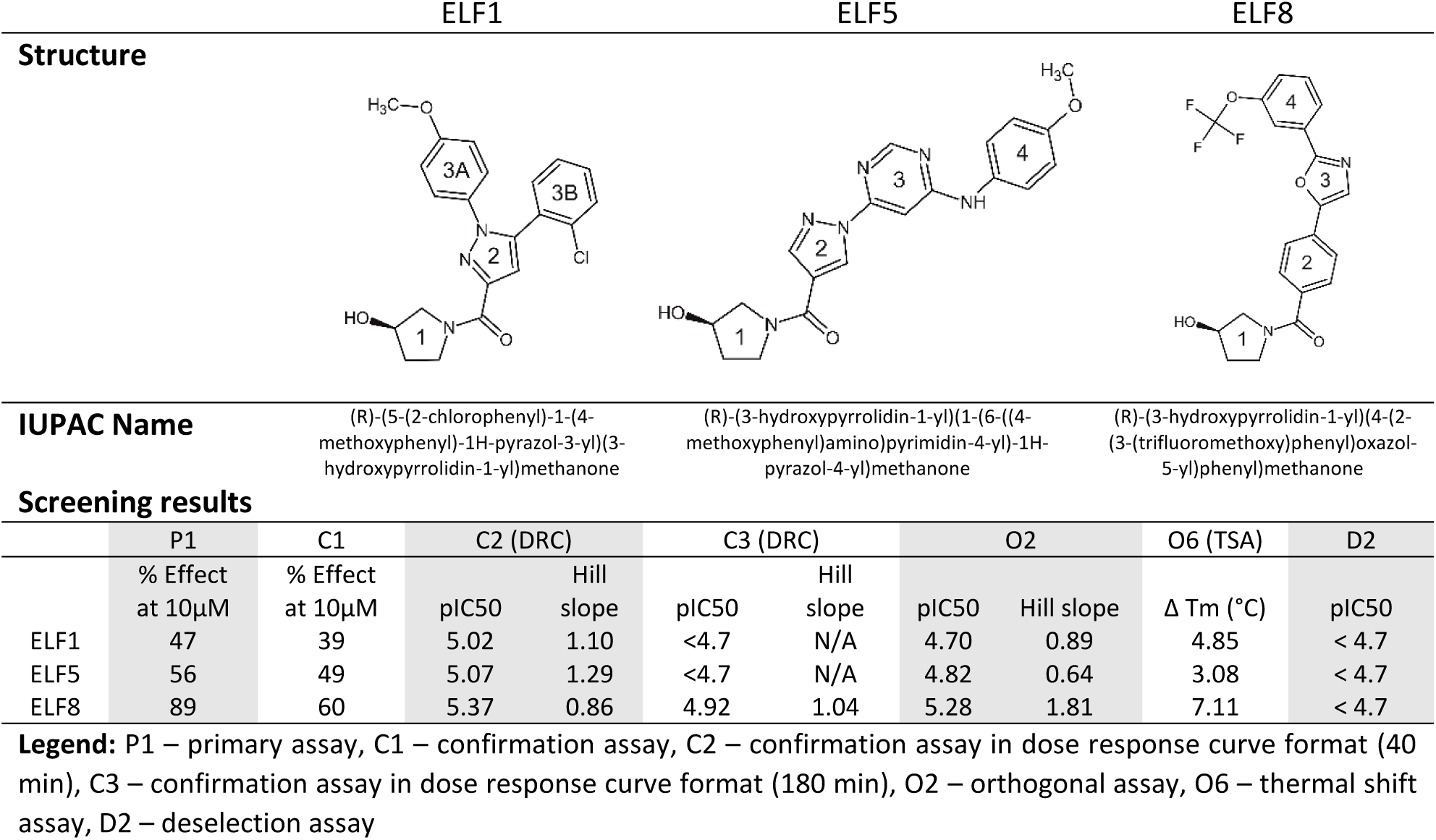
Summary of compound structures and screening results.

### Description of three selected Tier 1 compounds

In this paper, we describe the crystal structures we obtained for three of the Tier 1 compounds (Figure 2), termed ELF1 (*R*)-(5-(2-chlorophenyl)-1-(4-methoxyphenyl)-1*H*-pyrazol-3-yl)(3-hydroxypyrrolidin-1-yl)methanone, ELF5 (*R*)-(3-hydroxypyrrolidin-1-yl)(1-(6-((4-methoxyphenyl)amino)pyrimidin-4-yl)-1*H*-pyrazol-4-yl)methanone, and ELF8 (*R*)-(3-hydroxypyrrolidin-1-yl)(4-(2-(3-(trifluoromethoxy)phenyl)oxazol-5-yl)phenyl)methanone in complex with Rv0183. All compounds contain four ring systems (Table 2) A 3-hydroxypyrrolidin-1yl-methanone moiety as substituted heterocycle stands out as common element and novel chemotype for inhibition of MGL in all compounds. In this paper, we refer to the 3-hydroxypyorrolidin ring as ring 1. The second substitutions on the ketone are either a substituted pyrazol (ELF1, ELF5) or a substituted benzene ring (ELF8), each referred to as Ring 2. The pyrazole ring (ring 2) of ELF1 is substituted at two positions, namely at position 1 with a 4-methoxyphenyl group (ring 3A) and at position 5 with a 2-chlorophenyl ring (ring 3B). ELF5 and ELF8, on the other hand, are modified only at one position in ring 2, however, in each case with two additional ring systems. Ring 2 of ELF5 is a pyrazol that is substituted at positions 1 with a (methoxyphenyl)amino)pyrimidin-4-yl. For ELF5, we refer to the pyrimidin as ring 3 and to the methoxy-aniline as ring 4. In ELF8, ring 2 is a phenyl substituted in the para position with an oxazole (ring 3), which in turn is substituted with a (trifluoromethoxy)phenyl group (ring 4).

**Figure 2.**
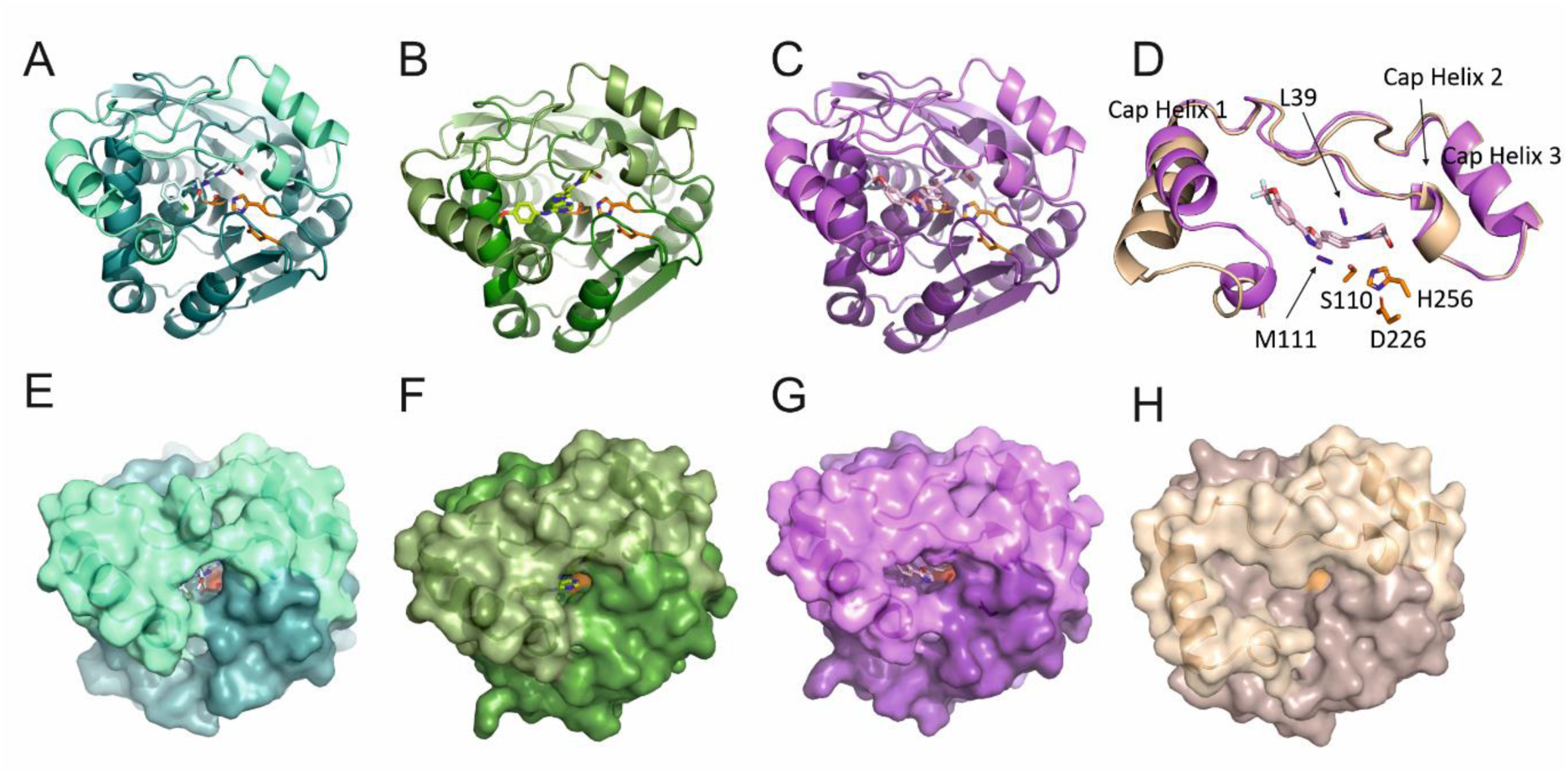
Overall structure of Rv0183 in complex with ELF inhibitors. The complex structures of the Rv0183 in complex with ELF1 (**A, E**, teal), with ELF5 (**B, F**, green) and ELF8 (**C, G**, violet) are shown in cartoon and surface representation. **(D)** Comparison of the cap region of Rv0183 in complex with ELF8 in almost closed conformation to the cap in open conformation as observed from a different experiment (PDB-ID: 6EIC, chain C^28^). **(H)** Surface representation of Rv0183 in open conformation (brown, PDB-ID: 6EIC). The α/β-hydrolase core is shown in darker colors, the cap region of each structure in lighter colors in all panels. Active site residues (S110, D226 and H256) are indicated as orange sticks.

All compounds have IC50 values in the single digit-µM range in the primary and orthogonal assay. The highest inhibition was observed for ELF8 (pIC50 = 5.37, IC50 = 4.26 µM), followed by ELF5 (pIC50 = 5.07, IC50 = 8.51 µM) and the branched compound ELF1 (pIC50 = 5.02 µM, IC50 = 9.55) in the primary assay using free glycerol as readout. The same trend was observed for the orthogonal assay using non-esterified fatty acids as readout confirmed this trend pIC50 (ELF8)> pIC50 (ELF5)> pIC50 (ELF1) (see Table 2). Due to the stringent selection process, the 3 compounds exhibited favorable physicochemical properties e.g., a molecular weight below 500 g/mol, LogD values below 5, calculated topological polar surface areas below 110 Å^2^, more H-bond acceptors than H-bond donors (HBA/HBD), and a Quantitative Estimate of Drug-likeness (QED) greater or equal than 0.67 (Table 3). ELF5 exhibits the highest total polar surface area (TPSA) value (105.4 Å^2^), indicating a higher degree of polarity, which could potentially impact membrane permeability. Additionally, ELF5 displays the lowest LogD value (1.56), indicating increased solubility in water. Furthermore, ELF5 exhibits the highest number of hydrogen bond acceptors and hydrogen-bond donors (HBA/HBD = 7/2). This is corroborated by the crystal structure, which reveals one additional bond compared to the other compounds. It has been established that the molecular weight (MW) of the compounds is very similar and within the optimal range for the development of the compounds. The QED values, a computed combination of molecular descriptors and properties associated with favorable drug-like characteristics are also within a narrow range, with ELF1 having the highest score (0.73), closely followed by ELF5 (0.69) and ELF8 (0.67).

**Table 3.** List of selected compound properties.

| Compound properties |  |  |  |  |  |
| --- | --- | --- | --- | --- | --- |
|  | TPSA | HBA/HBD | LogD | MW | QED |
| ELF1 | 67.59 | 4/1 | 3.73 | 397.85 | 0.73 |
| ELF5 | 105.4 | 7/2 | 1.56 | 380.40 | 0.69 |
| ELF8 | 75.8 | 4/1 | 4.37 | 418.37 | 0.67 |

**Legend:** TPSA - Topological Polar Surface Area, HBA/HBD – Hydrogen bond acceptor/donor, LogD – Distribution coefficient, MW – Molecular weight, QED - Quantitative Estimate of Drug-likeness.

### The overall structure of Rv0183 bound to inhibitors

We determined the crystal structure of Rv0183 in complex with ELF1, ELF5 and ELF8 at <2 Å resolution. The X-ray data collection and refinement statistics of the three complexes are presented in Table 4. The structure of Rv0183 (PDB-ID: 7P0Y^29^) was used for molecular replacement. In all structures, each asymmetric unit contains one molecule Rv0183 and one molecule inhibitor.

**Table 4.** X-ray collection and refinement statistics for Rv0183 inhibitor complexes. Values for the highest resolution shell are shown in parentheses. Data are generated using phenix.table_one.

| PDB-ID: | 9RHW | 9RF2 | 9RF5 |
| --- | --- | --- | --- |
| Inhibitor name | ELF1 | ELF5 | ELF8 |
| **Data collection** |  |  |  |
| Wavelength (Å) | 1.03272 | 1.03323 | 1.03323 |
| Space group | P 2 21 21 | P 2 21 21 | P 21 21 2 |
| **Cell dimensions** |  |  |  |
| *a*, *b*, *c* (Å) | 40.48, 82.53, 90.59 | 40.41, 82.78, 90.70 | 82.69, 91.10, 40.43 |
| $\alpha$, $\beta$, $\gamma$ (°) | 90, 90, 90 | 90, 90, 90 | 90, 90, 90 |
| Resolution (Å) | 45.30 - 1.25 (1.29 - 1.25) | 41.39 - 1.6 (1.66 - 1.60) | 37.65 - 1.8 (1.96 - 1.80) |
| *R*<sub>merge</sub> | 0.01947 (0.2044) | 0.01551 (0.4232) | 0.1049 (0.8358) |
| *I* / σ*I* | 15.92 (3.60) | 17.54 (1.65) | 21.21 (5.09) |
| Completeness (%) | 98.22 (86.69) | 99.49 (96.85) | 99.85 (99.96) |
| Redundancy | 2.0 (1.8) | 2.0 (1.9) | 12.8 (13.3) |
| CC ½ | 0.999 (0.857) | 1 (0.773) | 0.999 (0.931) |
| **Refinement** |  |  |  |
| No. reflections | 83267 (7253) | 40788 (3905) | 29038 (2844) |
| *R*<sub>work</sub> | 0.1554 (0.2076) | 0.1900 (0.3041) | 0.2254 (0.2497) |
| *R*<sub>free</sub> | 0.1647 (0.2184) | 0.2031 (0.3514) | 0.2520 (0.2730) |
| **No. atoms** | 2799 | 2336 | 2351 |
| Protein | 2428 | 2189 | 2211 |
| Ligand/ion | 94 | 85 | 59 |
| Water | 277 | 62 | 81 |
| **B-factors (Å<sup>2</sup>)** | 16.96 | 30.84 | 21.61 |
| Protein | 15.44 | 30.19 | 21.12 |
| Ligand/ion | 22.54 | 44.41 | 29.51 |
| Water | 28.33 | 35.14 | 29.25 |
| **R.m.s. deviations** |  |  |  |
| Bond lengths (Å) | 0.014 | 0.005 | 0.007 |
| Bond angles (°) | 2.15 | 0.80 | 0.91 |
| **Ramachandran** |  |  |  |
| Favored (%) | 98.19 | 97.83 | 98.19 |
| Allowed (%) | 1.81 | 2.17 | 1.81 |
| Outliers (%) | 0.00 | 0.00 | 0.00 |
| Rotamer outliers (%) | 0.39 | 2.61 | 3.02 |
| Clashscore | 10.89 | 4.02 | 8.82 |

The overall structure of the protein shows an 8-stranded central, twisted β-sheet that is flanked on both sides by the connecting helices (Figure 2, Panels A-C). The cap (A138-P193) is an extension of a loop between β-strand 6 and the flanking helix connecting the cap to strand 7. The cap covers the substrate binding cavity and has a Z-like shape that is formed predominately by three α-helixes for Rv0183 (Figure 2, panels A-D). In this manuscript, we term these short helices cap helix 1, cap helix 2, and cap helix 3 (Figure 2, panel D).

The active site residues are arranged to form a catalytic triad: The nucleophile S110 is in the GXSXG motif of the nucleophilic elbow directly after β-strand 5. D226 is in a loop after β-strand 7 and H256 is located in a loop region connecting β-strand 8 with the C-terminal helix. The backbone atoms of residues M111 and L39 are in position to form the oxyanion hole (Figure 2). All inhibitors are bound in the substrate binding pocket and are very close to the active site, yet do not show covalent binding modes (Figure 2, Panels A-D).

Conformational flexibility of the cap affects the shape and solvent accessibility of the substrate binding pocket. All three Rv0183-ELF-complex structures presented here display an almost entirely closed conformation (Figure 2, panels E-G). Conformational dynamics must have taken place for the inhibitors to bind within the observed almost closed cavity. Cap helix 1 (V146-V157) has a significant impact on the shape of the binding pocket and has been described as primarily responsible for constraining access to the pocket. In the closed conformation as observed here, cap helix 1 tilts towards the entrance of the binding pocket (Figure 2D), as also shown previously for a complexed structure of Rv0183 (PDB: 7OZM, ^29^). For comparison of the large conformational flexibility, full access to the active site in an open conformation was observed previously for an uncomplexed structure of Rv0183 (Figure 2H, PDB-ID: 6EIC, chain C, ^28^).

Although all ELF-complex structures adopt a highly similar, almost closed cap conformation (RMSD of the cap residues are 0.24 Å (Rv0183-ELF5 to Rv0183-ELF1) and 0.44 Å (Rv0183-ELF8 Rv0183-ELF1) and 0.29 Å (Rv0183-ELF5 to Rv0183-ELF8), individual residues or small stretches of the cap show high B-factors and different side-chain orientations (Figure 3).

**Figure 3.**
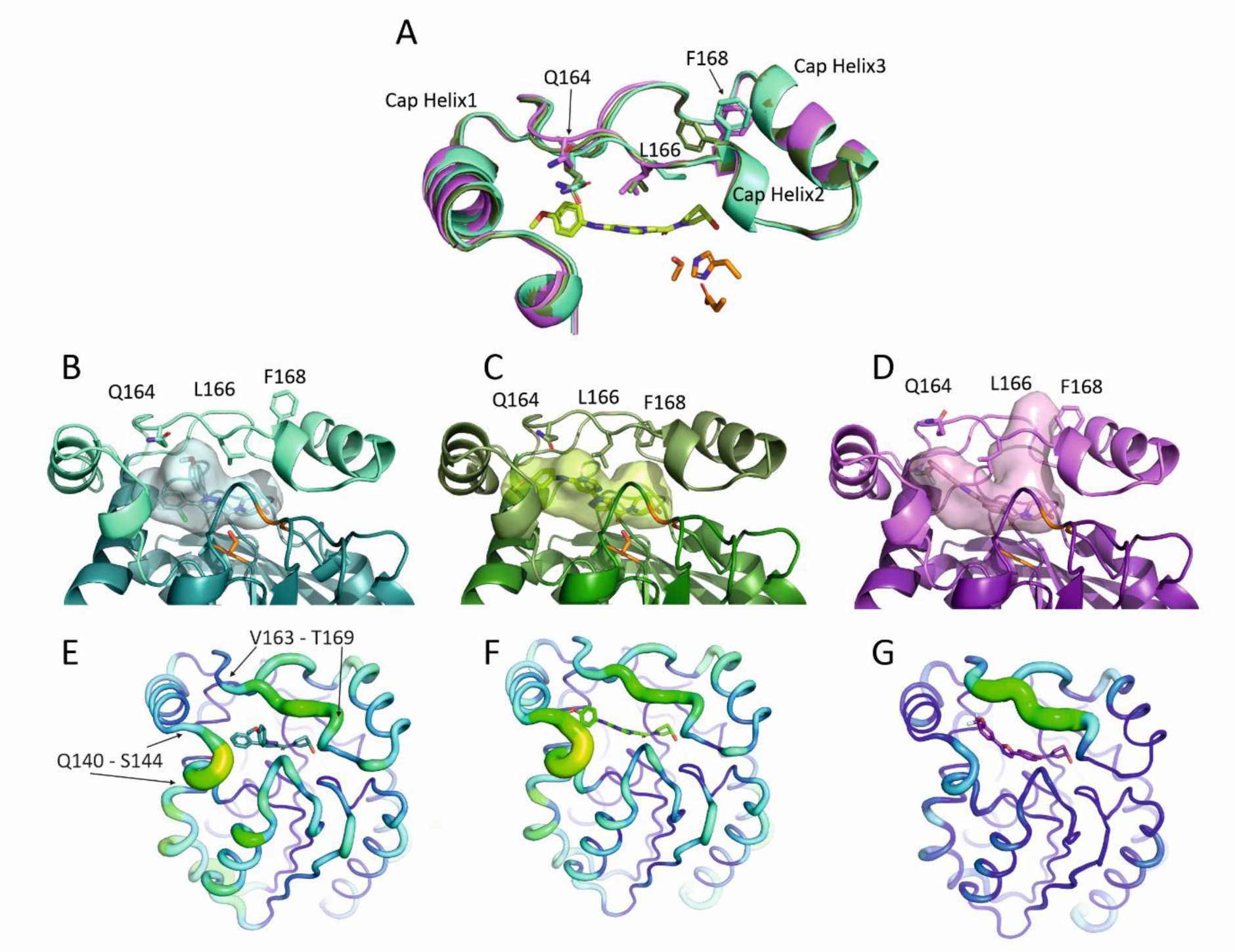
Conformational flexibility and dynamics the cap region. **(A)** Dynamics of specific cap residues shape the binding pocket. Overlay of all three structures as cartoon with ELF5 and specific flexible residues highlighted as sticks. **(E-G)** B-factor representation in complex structures. The complex structures of the Rv0183 in complex with ELF1 (**B, E**, teal), with ELF5 (**C, F,** green) and ELF8 (**D, G**, violet) are shown in cartoon and surface representation, the cap-structures are in lighter color-variants. Active site residues (S110, D226 and H256) are indicated as orange sticks.

Such high B-factors indicating high conformational mobility are observed in the loop region from Q140 to S144 and V163 to T169 residue. Especially movement of Q164, L166 and F168 strongly influence the shape of the substrate/inhibitor binding pocket and its accessibility (Figure 3E-G).

Residues L166 and F196 are also observed to adopt different conformations leading to the opening of the binding pocket between the short cap helix 2 (F168-I171) and cap helix 3 (P175-T183). In the Rv0183 structure complexed with ELF1, L166 restricts the binding pocket by pointing inwards, towards the core of the protein (Figure 3A-B), whereas in the ELF5-complex structure F168 is responsible for restricted access (Figure 3C-D). In the ELF8-Rv0183 structure, side chains of L166 and F168 point towards the surface, enabling a small, second access channel to the active site. In the structure of Rv0183 with ELF5, another residue with a higher B-value, Q164, moves towards the inhibitor and stabilizes inhibitor binding with an additional hydrogen bond in ELF5-Rv0183 (Figure 3A, 3C, Figure 5E).

Different parts of the substrate binding cavity of Rv0183 have different electrostatic properties (Figure 4A-C). The entrance is mostly hydrophobic, whereas the area in close proximity of the catalytic triad is polar. We observe that the possible second access channel between cap helix 2 and cap helix 3 is lined by main-chain carbonyl-oxygens pointing towards the binding pocket and by side-chains with negative charges e.g., E41, E165, D167, E256. At least one water molecule was assigned to this channel. Another small pocket is separated from the negatively charged part of the binding cavity by a salt bridge formed between R45 and E257 (distances (N-CO) of 2.9 Å). Electron density in the small cavity has also been attributed to water molecules (Figure 4D-F). E257 is additionally involved in binding of the OH-group of ring 1 in all three inhibitor-Rv0183 structures presented here (see below, Figures 5).

**Figure 4.**
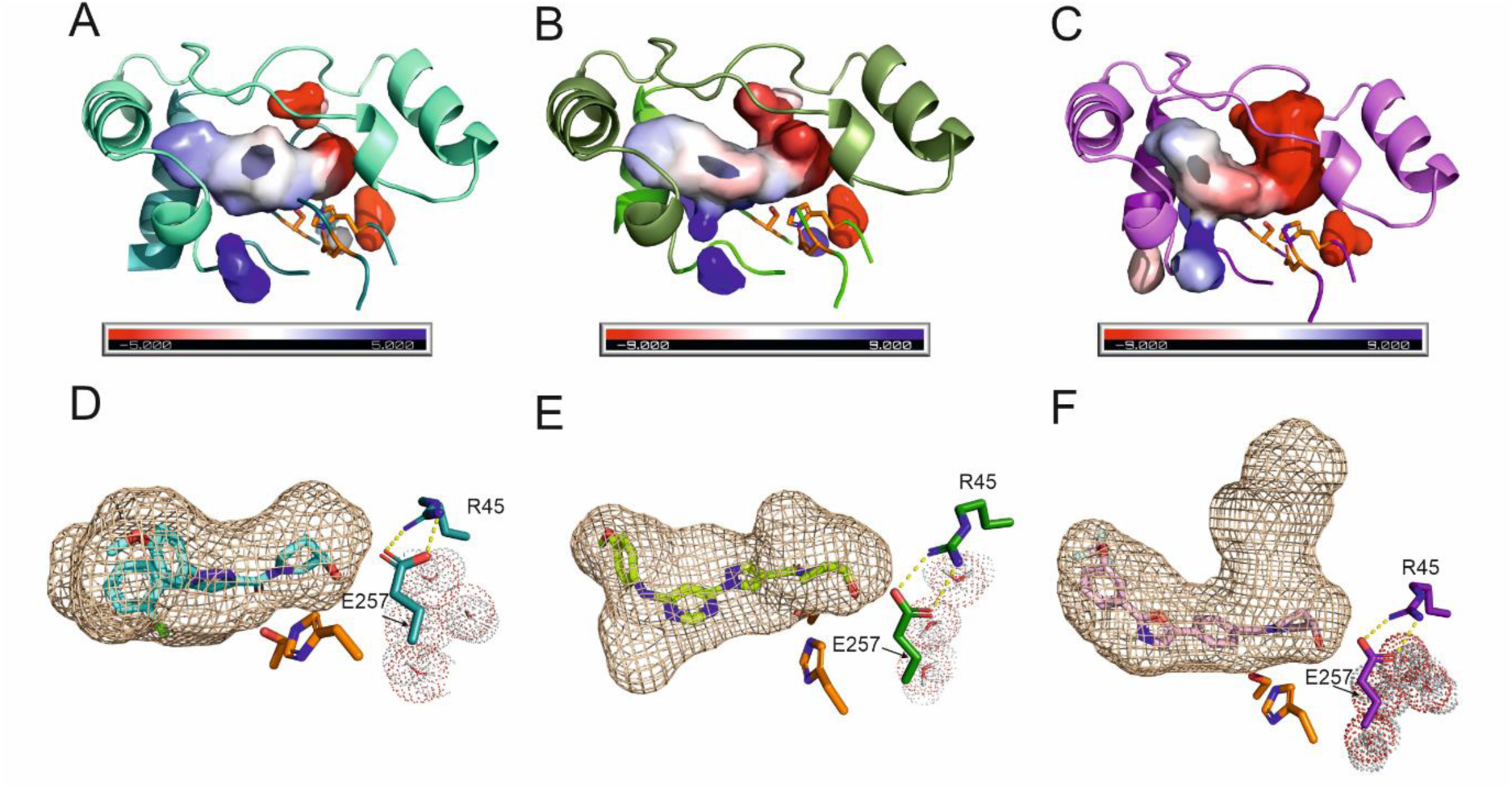
Electrostatics and general shape of the cavity. **(A-C)** Different parts of the cavity of Rv0183 in complex with ELF1 (**A**, cyan), ELF5 (**B**, green), and ELF8 (**C**, pink) have different electrostatic properties. Active site residues as orange sticks. **(D-F)** Cavity of Rv0183 with inhibitors ((**D)** ELF1 in cyan, (**E**) ELF5 in lime, (**F**) ELF8 in pink) bound is separated from small water-filled cavity by salt-bridge built by R45 and E257 (stick representation). Distance measured is 2.9 Å. Active side residues S110 and H256, are shown as orange sticks. Some residues are omitted for clarity.

**Figure 5.**
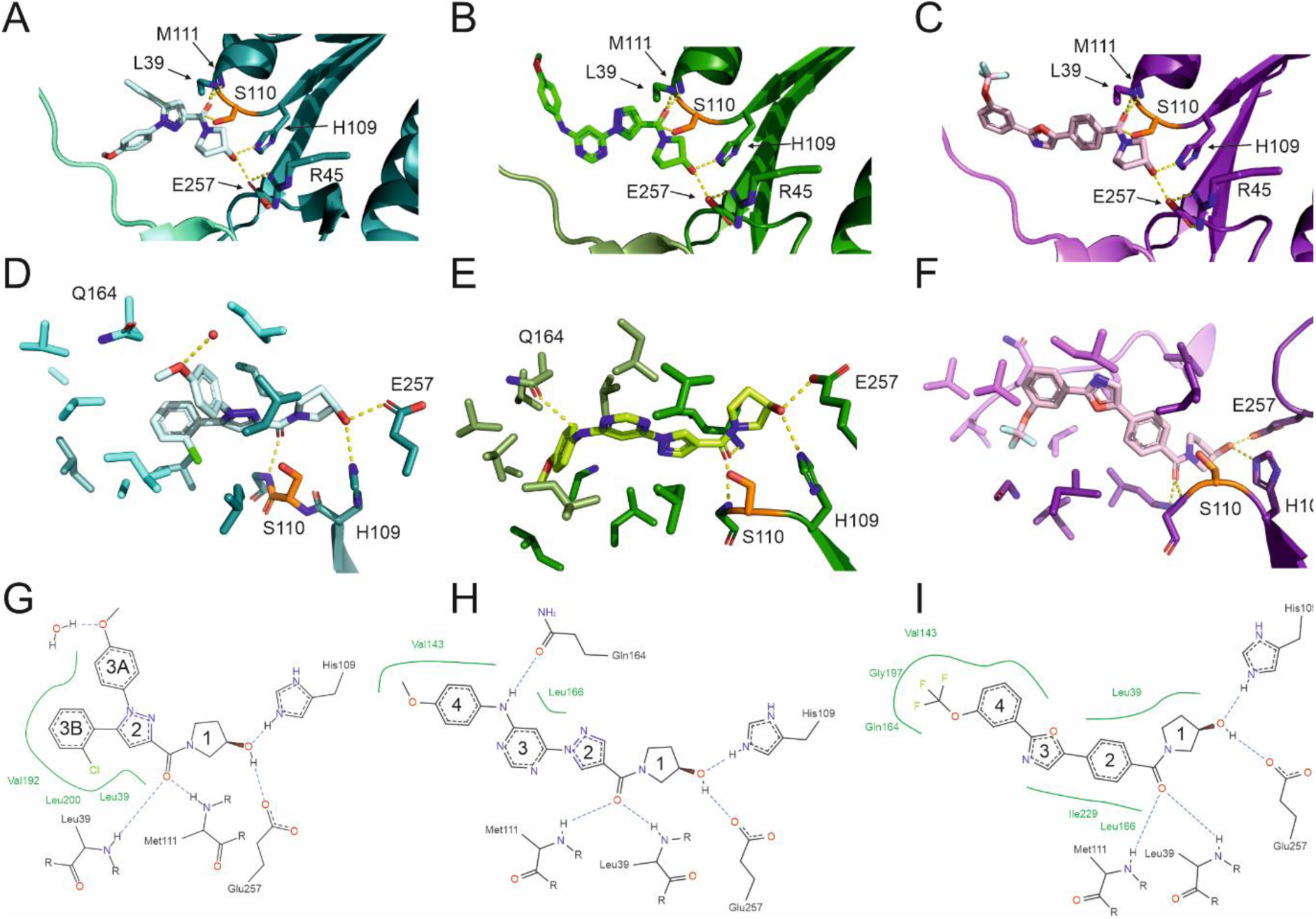
Details of ELF-ligand interaction with Rv0183 as observed in the high-resolution crystal structures. (A-F) The complex structures of the Rv0183 in complex with ELF1 (**A, D**, in teal, left panel), with ELF5 (**B, E**, green-shades) and ELF8 (**C, F**, magenta shades) are shown in cartoon and stick representation, the cap-structures are in lighter color-variants. Cavity lining and H-bonding residues are depicted as sticks. Active site residues (S110, D226 and H256) are indicated as orange sticks. **(G-I)** 2D view of the inhibitors in the binding pocket **((G)** ELF1, **(H)** ELF5, **(I)** ELF8)

### Specific interactions of the ELF inhibitors with Rv0183 facilitate efficient binding

ELF1, ELF5, and ELF8 consists of 4 ring systems (Table 2, Figure 5). Ring 1 and the adjacent keto group are bound in the same conformation in all complex structures (Figure 5). The oxygen of S110 is 2.8-2.9 Å away from the carbonyl-carbon of ELF1, ELF5 and ELF8 and the distances of carbonyl-oxygen connecting ring 1 and ring 2 from the inhibitor to main-chain nitrogens of the oxyanion-hole (L39, M111) are 2.7 - 2.9 Å for all three inhibitors (Figure 2D and Figure 5D-I). The OH-group of ring 1 forms a H-bond network with H109 and E257 of the lipase core (Figure 5D-I). Ring 2 is either a substituted pyrazole (ELF1, ELF5) or a substituted benzene ring (ELF8) and still adopt similar positions in the complex structures (Figure 5).

Detailed analysis of ELF1 binding (Figure 5A, 5D, 5G) reveals that the compound is embedded by residues L39, H109, S110, M111, L142, V143, Q164, L166, V192, L200, L228, I229, E257. In addition to the H-bond network that was previously described, a hydrogen bond between a water molecule and the ligand is observed for the oxygen atom of the methoxy substituent on ring 3A. This atom functions as a hydrogen bond acceptor. The side chain of L39 is located underneath the aromatic pyrazol of ELF1 (ring 2) in a distance less than 5Å. The phenyl-group of ring 3A is embedded within a cavity lined by the hydrophobic side chains L142, L166, V192, L228, I229. The methyl-group of ring 3A reaches the surface of Rv0183 between the core and cap helices 1 and 2. (Figure 2A, 2E, Figure 5). The chlorophenyl-ring (ring 3B) also branching off from ring 2 stays deeply within the binding pocket and is embedded within the hydrophobic residues L39, L142, V143, V192, I196, L200, I229, and the aliphatic parts of the side chain of Q164.

In the structure with the inhibitor ELF5 rings 1 and 2 occupy the same region of the binding cavity like observed in the structure with inhibitor ELF1. The H-bond network between the protein and ring 1 of ELF5 is the same as for ELF1 (and ELF8) (Figure 5). Side-chains of L39, L166, L200, L228 and I229 line the cavity within 5Å to the pyrazole ring (ring 2). Parts of the pyrimidine (ring 3) of ELF5 can be seen via the entrance tunnel in the complex structure, in a similar position to ring 3A of ELF1 (Figure 2B, 2F, Figure 5). Ring 3 of ELF5 is placed in a hydrophobic pocket lined by side-chains of L142, L166, L200, L228 and I229 (within 5 Å). The side chain methyl groups of L228 are building a hydrophobic environment for ring 1, 2 and 3 (3.9-5.0 Å). The NH-group connecting the pyrimidine ring (ring 3) to the methoxyphenyl-ring (ring 4) acts as H-bond donor to the side-chain carbonyl oxygen of Q164, adding an additional H-bond to the protein-inhibitor interaction. The phenyl-ring (ring 4) of ELF5 then proceeds within the binding cavity enclosed by cap residues, lined by side-chains of A139, L142, V143, V147, V192, L200, and the aliphatic part of Q164. The methyl groups of V143, V147, V151 and L201, along with the C-alpha of G197 form a hydrophobic cavity (within 4 Å radius) well suited for the distal methylgroup of ring 4.

The binding of ring 1 in ELF8 is identical to those described before (Figure 5C, 5F, 5I). Ring 2 of ELF8 is located in a hydrophobic pocket lined by the side chains of L39, L166, L200, L228 and I229. The oxazole (ring 3) of ELF8 is embedded in a central part of the cavity lined by L142, L166, L200, and I229 from the cap and the core of Rv0183. The (trifluoromethoxy)phenyl ring (ring 4) of ELF8 is positioned in a hydrophobic pocket primarily formed by cap residues V143, V147, A151, V163, G164, V192, and G197. Thus, the methoxy-group is neatly buried underneath the cap.

### In silico characterization of Rv0183 ligand interactions and selectivity

#### Molecular docking validates strong binding of ELF8 and variation in binding modes of ELF1 and ELF5

To validate the experimental binding affinity results and assess the suitability of the system for future *in-silico* inhibitor refinement pipelines, molecular docking was used to predict the binding affinities of ELF1, ELF5, and ELF8 to available crystal structures of Rv0183 (Figure 6). Analyzing the lowest predicted binding energies from 45 independent docking runs, ELF8 exhibited the strongest binding to all six crystal structures, with an average binding energy of -12.1 ± 0.5 kcal/mol. ELF5 showed an average binding energy of -10.3 ± 0.9 kcal/mol, while ELF1 binding energy averaging -9.9 ± 0.3 kcal/mol (Figure 6). Notably, the lowest-energy binding mode of ELF1 very closely matched the experimentally determined conformation in overall orientation and positioning. Nevertheless, ELF1 displayed varying conformations in its lowest-energy predicted binding modes with different orientations and positioning of the ring systems. ELF5’s binding mode was consistent in four out of six target structures, reflecting the orientation and general positioning of the ring system as experimentally determined. The general positioning of the ring-system of ELF8 in the binding pocket was resembled by all top energies across six different crystal structures of Rv0183, indicating robust and specific binding, and demonstrating high suitability for in silico engineering.

**Figure 6.**
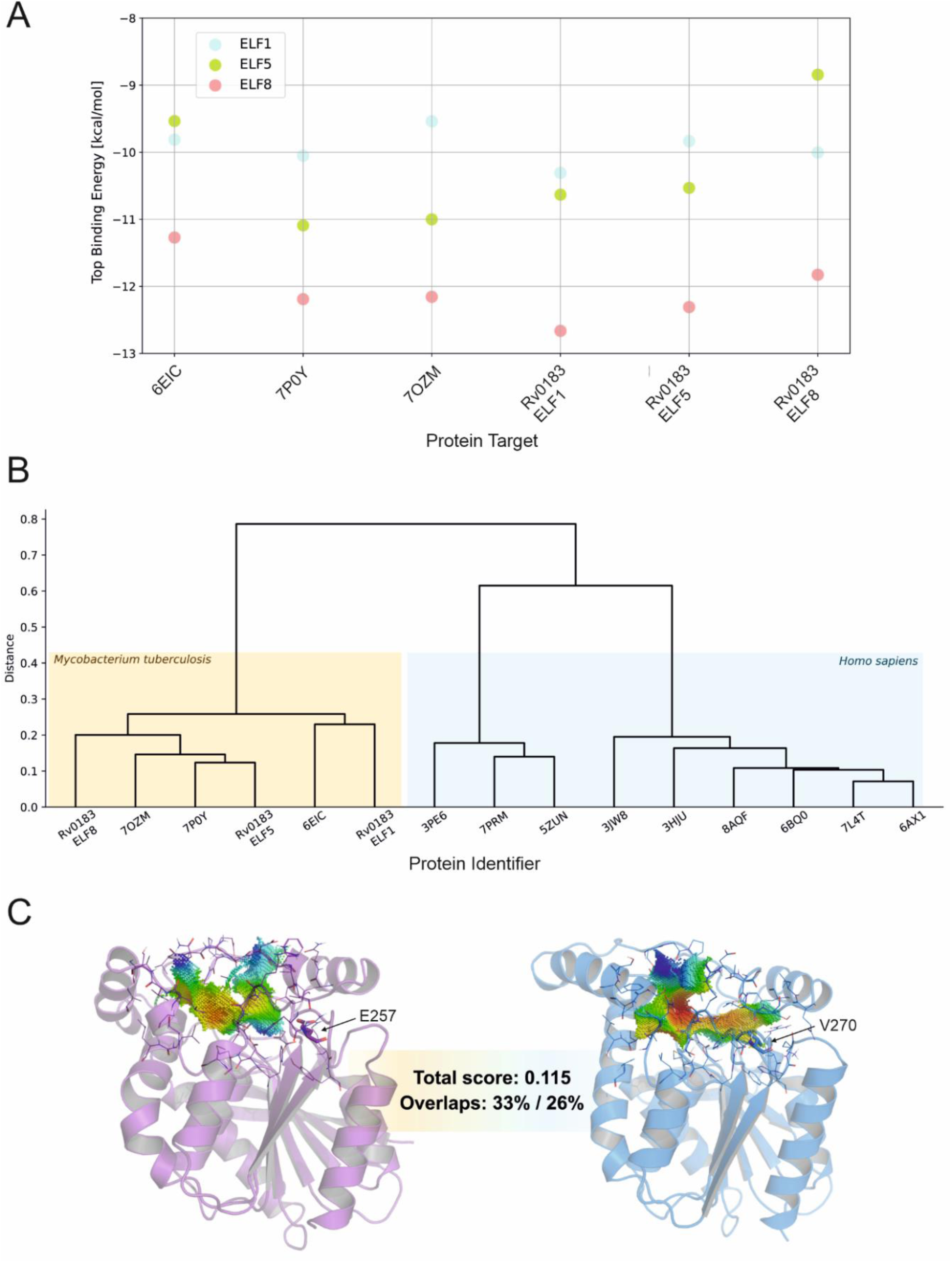
In silico studies. (A) Molecular docking of ELF-compounds into Rv0183. Binding Energies of ELF1 (cyan), ELF5 (green) and ELF8 (pink) determined via docking into cavities calculated from available Rv0183 structures ^28,29^. **(B)** Comparative analysis of binding sites in human orthologs. Distance tree for point-clouds calculated from experimentally determined cavities of human MGL ^52–60^ (blue background) and Rv0183 (yellow background). **(C)** Cavities of human MGL (7PRM ^53^, blue) compared to that of Rv0183-ELF8 (pink) highlighting the difference of the polar E257 in Rv0183 compared to the aliphatic V270 in human MGL. E257 forms a salt-bridge delineating the binding cavity of Rv0183 and H-bonds with the 3-hydroxypyorrolidin ring common in all three identified inhibitors. Point clouds colored by hydrophobicity (hydrophobic to hydrophilic: red to blue)

#### Comparative analysis of binding sites in human orthologs emphasizes the potential for selective inhibition of Rv0183

MG-hydrolyzing activity in humans is primarily attributed to human MGL (hMGL, experimental structures are known, e.g., PDB 7PRM, (UniProt Accession code: Q99685) ^61^. To a much lesser extent, physiologically relevant MG hydrolytic activities are attributed to ABHD6 (UniProt Accession code: Q9BV23; PDB-code: 7OTS^62,63^), and ABHD12, for which no experimental 3D structure is available (UniProt Accession code Q8N2K0). A pairwise comparison of inhibitor-binding sites between available structures of the mycobacterial MGL Rv0183 and those of the human MGL orthologs reveals that Rv0183s form a distinct cluster, separate from the human MGL ortholog (Figure 6B). In this analysis, the open conformation of Rv0183 (6EIC) forms a separate branch within the mycobacterial proteins. While a qualitative comparison of cavities is difficult, the CatalaphoreTM-methods also gives quantitative scores for cavity matches. A lower matching score indicates greater cavity similarity, with identical cavities achieving a total score of 0. Another metric for cavity similarity is the overlap of the query and target cavities. These values, expressed as percentages, indicate the volume of each respective cavity that matches with the compared cavity. In the direct comparison of Rv0183 (as present in the ELF8 complex) and human MGL (7PRM; 36.7% sequence identity) we achieve a total score of 0.115 and volumetric overlaps of 33%/26% (Figure 6C). The matched portion corresponds to 33% of the 7PRM cavity. 26% represents the proportion of the Rv0183 cavity that overlaps with the human MGL cavity following alignment of the 3D point cloud. Consequently, the total score is calculated based solely on this overlapping region. Human ABHD6 (18% sequence identity with Rv0183) also has an α/β-hydrolase structure with a somewhat similar cavity shape, leading to a total score of 0.187 and 32%/25% overlap. Human ABHD12 (26% sequence identity with Rv0183) has a predicted similar overall structure, yet the cavities show very little overlap preventing meaningful cavity matching (at least without additionally integrating dynamics of cavity-lining residues).

This analysis, based on physico-chemical properties and cavity shape—both critical factors for ligand binding—suggest that there is great potential for designing inhibitors that specifically target Rv0183. Such specificity could lead to a favorable toxicity profile, allowing the development of effective and save therapeutics.

### Whole cell testing of compounds in Mtb

Based on prior transposon mutagenesis and genome-wide CRISPRi studies, *rv0183* has been characterized as a non-essential gene for growth of Mtb in standard growth medium ^25,64^, likely due to high concentrations of glycolytic (glucose and glycerol) carbon sources that could bypass the requirement of a monoacyl glycerol lipase for growth. We therefore reasoned that testing the effects of the lead compounds for Mtb growth inhibitory activity would require a more relevant medium composition in which growth is dependent on monoolein catabolism. To this end, we designed a synthetic growth medium in which monoolein was the sole available carbon source (see materials and methods) and could show that *Mtb*H37Rv grew normally (similar doubling time to standard 7H9-based, glucose/glycerol-containing medium) under these conditions (data not shown). We then conducted microtiter broth dilution MIC assays of the 3 lead compounds against *Mtb*H37Rv in this medium. As shown in Figure 7, all 3 compounds were able to inhibit the growth of *Mtb*H37Rv, albeit at high micromolar concentrations (MIC values > 50 µM).

**Figure 7.**
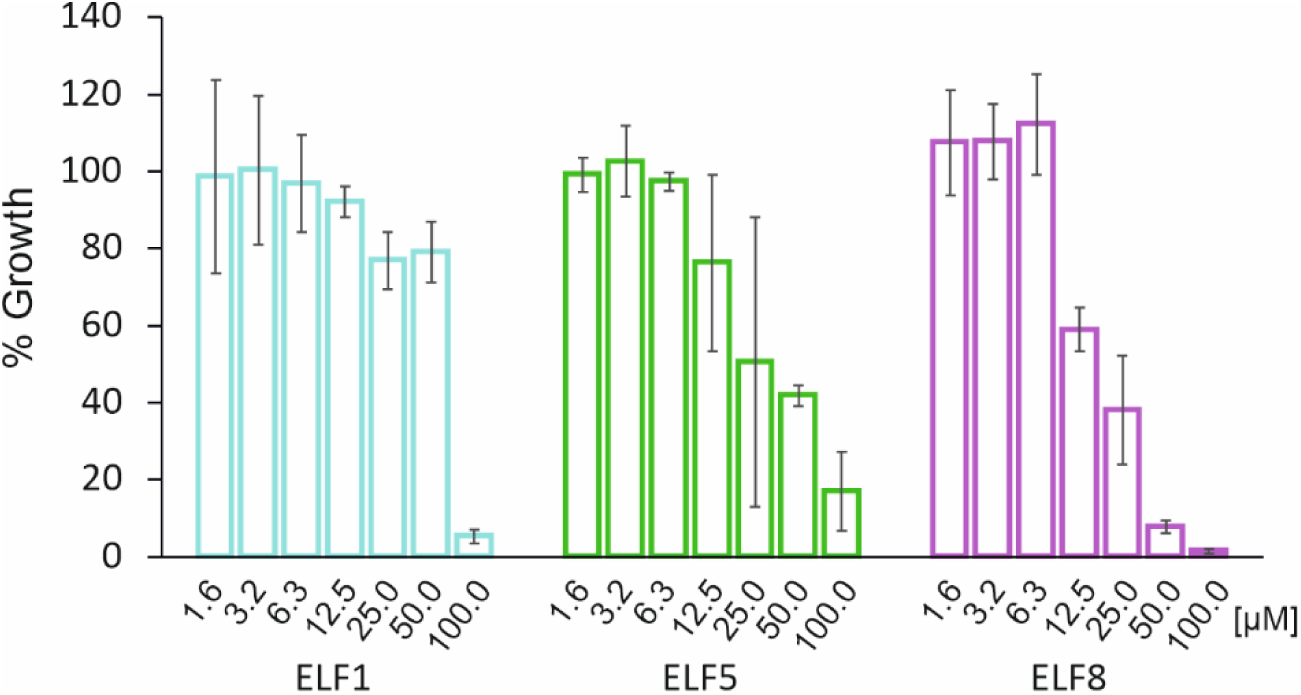
Whole cell testing on synthethic growth medium with monoolein as sole available carbon source. All compounds were able to inhibit the growth of MtbH37Rv at 100µM compound concentration.

## Discussion

In this research, we describe the identification and detailed structural characterization of 3 compounds out of a library of 470,000 that act as potent inhibitors for the lipase Rv0183. The inhibitors have a low MW (398-418 Da) and good druggability scores, IC50 values (4.2-9.5µM) and share a hydroxypyrrolidine as a common element that is bound close to the active site. These inhibitors represent a valuble group of potential compounds for selective inhibition of Rv0183 upon for further development.

Different structures of Rv0183 have been published, both in free and substrate-mimic-bound forms and demonstrate a large substrate binding pocket that is formed by the core lipase and the cap thus fine-tuning the overall shape of the cavity ^28,29^. Similar cap dynamics has been observed structurally for other monoacylglycerol lipase, namely human MGL, MGL from *Bacillus* sp. H257 and the yeast monoacylglycerol lipase YJU3p ^65^. Although the volume of the binding pocket varies due to its dynamic nature, the physicochemical properties remain consistent.

In the three structures published here, the residues of the second helix of the cap, namely Q164, E165, L166, and F168 seem to have high flexibility with respect to side-chain placement either inwards towards the substrate binding cavity or towards the surface of the protein. Side-chain flexibility in this part of the cap has also been observed for orthologs of Rv0183, with Leu166 of Rv0183 corresponding to the so-called gatekeeper residue I145 from *Bacillus sp*. H257 and I178 in the human ortholog ^33,66^. Flexibility of the cap can open up another entrance to the binding pocket. It has been proposed for other MGL homologs from human and *Bacillus* sp. H257 that the glycerol product of the enzymatic reaction can exit the binding pocket through a glycerol exit channel ^66,67^. Some density indicating an alternate conformation for ring 1 with the OH group pointing to the opposite direction is also present in the complex structure with ELF5. In this conformation, a weak H-bond might form between the OH-group and Y181 of Rv0183 (3.5 Å O-O distance, not shown) that is reminiscent of the glycerol-binding by E256 in MGL from *Bacillus* sp. H257.

The ELF inhibitors create a network of hydrogen bonds around the active site to the hydroxypyrrolidine, while the other ring systems are located within a cavity mainly formed by hydrophobic side chains. These inhibitors are positioned in the binding pocket similarly to the substrate mimic maglipan (hexadecylbutylphosphonofluoridate)^29^. The hydroxypyrrolidine adopts a position similar to the glycerol moiety in the structure of MGL from *Bacillus sp*. H257 (variant D196N for catalytic inactivity) in complex with the substrate 1-laureoyl-glycerol ^33^. For ELF1, ELF5 and ELF8 the hydroxy-group of the hydroxypyrrolidine ring 1 is hydrogen-bonded to H109 and E257. At the hydrophilic bottom of the binding pocket of Rv0183, the nucleophile S110 can perform the nucleophilic attack on the carbonyl carbon, while the oxyanion is stabilized by the oxyanion hole residues, as described by Grininger et. al ^29^. In the ELF-inhibitor complexed structures, backbone NH-atoms of M111 and L39 are in hydrogen bond distance to the carbonyl-carbon at the equivalent position. The other parts of the ELF-inhibitors have mostly hydrophobic interactions in the tunnel formed by hydrophobic side chains. This portion of the pocket provides a favorable environment for the alkyl-chain of the monoacylglycerol substrate.

ELF1 differs prominently from ELF5 and ELF8 with respect to the branching that occurs at the pyrazole ring (ring 2). Therefore, ELF1 is shorter in length despite the same number of rings, but broader and the methoxy-phenyl (ring 3A) reaches the entrance to the surface of the protein. ELF8 and especially ELF5 are covered underneath the cap due to the elongated arrangement of the ring systems. ELF5 is the only inhibitor that has an additional hydrogen bond to the protein apart from ring 1, which is between the HN-connection between ring 4 and 5 and to the flexible cap residue Q164 (B-factor: 46.2 Å^2^).

ELF8 showed the best inhibition properties (pIC50 = 5.37) along with the highest increase in melting temperature (Δ Tm = 7.3 °C). The higher molecular weight (418 Da) and LogD (4.37) values is due its trifluoralkoxy-substitution on ring 4. Substitution of a methyl or alkoxy groups with a fluorinated homologues (e.g., trifluoromethyl (CF_3_) or OCH_3_ to OCF_3_ group) is used frequently in medicinal chemistry ^61^. The C-F bond has high strength, a large dipole moment, strong electronegativity, and small size. The increase of lipophilicity of a compound harboring a CF_3_ group often leads to advantageous modulation of biophysical and pharmacokinetic properties like hydrophobicity, electrophilicity and ligand-protein interaction (increase in logD, lower passive permeability). The -OCF_3_ group is sometimes also associated with increased metabolic stability, possible due to reduced O-demethylation and reduced oxidative metabolism ^68^. Considering further development of Rv0183 inhibitors, it its therefore very good to see that such advantageous trifluoromethoxy-benzene groups can smoothly be embedded in the ligand binding cavity of the target protein Rv0183. The binding poses of the ELF-inhibitors as observed experimentally were also identified by molecular docking within the CavitOmix platform with especially robust conformations and strongest predicted binding for ELF8 (see Figure 6A). This provides an outstanding starting point for further compound development aided by *in silico* approaches.

In the initial phase of the high-throughput screening one goal was to find species-selective inhibitors. In the deselection campaign we were looking for an inhibitor that did not bind to the structurally highly similar human homolog. Even though this criterion was not considered in the selection process initially, no compound in the QHL showed distinct activity on the human ortholog, demonstrating the strength and selectivity of the screening process.

Inhibition of human MGL has been a long-standing pharmacological target for neurodegenerative diseases, inflammation and cancer. The production of fatty acids by human MGL acts as pro-tumorigenic signal and fatty acids are used for tumor growth. The endocannabinoid 2-arachidonoylglycerol (2-AG) is a monoacylglycerol and a prominent metabolic target of human MGL in the brain. Raising endocannabinoid-levels via blockage of its metabolism, is a sought-after therapeutic option especially for chemotherapy-induced neuropathy and further nourished multiple drug-development campaigns ^61,69–75^. The goal of our study is to inhibit Rv0183 specifically, without affecting the human ortholog. This is important because of the observed negative effects of the efficient irreversible binding of general human MGL inhibitors, such as JZL-184, that cross the blood-brain barrier ^76^.

The sequence identity between human MGL and Rv0183 is conserved to 34%, although the overall structure is very similar. Jiang et al. recently reported the structure of a potent selective inhibitor, LEI-515, developed through high throughput screening against human MGL^57,74^ . The compound binds to human MGL through a covalent reversible mechanism (PDB-code: 8AQF). A comparison of the LEI-515 sulfonyldifluoropentanol moiety binding to human MGL with the ELF hydroxypyrrolidin binding to Rv0183 reveals similar main-chain mediated oxyanion hole hydrogen bond network to the carbonyl oxygen at the bottom of the binding pocket. However, the hydrophobic fluorine moiety of LEI-515 engages in strong hydrophobic interactions via V270 in human MGL, while on the equivalent position in Rv0183 the charged residue E257 plays a crucial role in expanding the hydrogen bond network with the hydroxypyrrolidin of the ELF inhibitor structures (Figure 6C). Thus, the HTS of more than 200,000 compounds selectively for human MGL did not lead to any structure containing a hydroxypyrrolidin in their QHL containing 50 compounds ^74^.

To investigate the differences in the binding pockets in more detail, we used a novel structural bioinformatics method described as the Catalophore^TM^ approach. This technology analyses bindingsite similarities by comparing 3D point clouds representing different physico-chemical properties projected by the amino acids into the cavity’s empty space. It allows the identification of similar binding sites independently of overall amino acid sequence identity or protein structure, thereby broadening the search space for structurally unrelated proteins which might still comprise similar function ^77^. Originally showcased to identify promiscuous biocatalytic activity of enzymes, the method has developed into an efficient tool for *in silico* drug-repurposing and side-effect screens ^43,78^. The different clustering of active site cavities from human proteins and Rv0183 further confirm great potential for designing inhibitors that specifically target the mycobacterial lipase.

Testing the ability of these compounds to inhibit Rv0183 at the whole cell level is complicated due to the non-essential nature of this gene under standard, aerobic growth conditions ^25,64^. However, by modifying the growth medium to make Mtb reliant on monoolein catabolism for growth, we were able to use standard microtiter plate MIC methodologies to demonstrate genuine growth inhibitory activity of the compounds against the bacterium. While these are promising preliminary results indicating potential on-target activity at the whole cell level, we have not assessed the growth-inhibitory activity of the same compounds with different carbon sources and therefore cannot explicitly conclude that compound activity is monoolein-specific. Furthermore, the MIC values of the 3 tested compounds (50-100 µM) were far in excess of the IC50s determined against purified recombinant protein; this could be due to cell permeability issues, but could also point to a mechanism of action independent of Rv0183. The role of Rv0183 in allowing Mtb to grow on monoolein as a sole carbon source would be best assessed using gene knockout/knockdown strains, however this is beyond the scope of this work.

To the best of our knowledge, no studies have investigated the role of monoacylglycerol metabolism on the physiology and life cycle of Mtb. It is unlikely Mtb relies on monoacylglycerol as a sole carbon source at any point during the infection cycle, due to the wide variety of carbon sources available within the host milieu ^79^. However, monoacylglycerol metabolism is likely to be an important process for lipid recycling (e.g., phospholipid metabolism and cell membrane remodeling), as well as an intermediate process in the catabolism of more complex lipid species, such as triacylglycerols, which are known to be carbon sources for the bacterium during host infection ^14,80,81^. Intracellular triacylglycerols accumulation and subsequent hydrolysis is also known to be important for *Mtb* to tolerate certain *in vitro* generated stress conditions, such as hypoxia ^82^ and nutrient starvation ^81,83^. It is therefore possible that Rv0183 inhibition would have a more dramatic effect under more infection-relevant conditions (such as growth within macrophages, or in *in vitro* non-replicating model systems), and therefore such assays would form the basis of any future microbiological work on these compounds.

## Conclusion

In conclusion, our study has made a significant advancement in the development of selective inhibitors against Mtb by targeting the lipase Rv0183. Guided by an extensive high-throughput screening campaign of nearly half a million compounds, we have identified a trio of hydroxypyrrolidine-ring compounds—ELF1, ELF5, and ELF8—as potent inhibitors, demonstrating species-specific interaction with their bacterial target. These inhibitors exhibit not only low micromolar IC50 values but also desirable drug-like physicochemical attributes, affirming their promise for further drug development. Structural elucidation and molecular docking have revealed their non-covalent binding within the active site of Rv0183 and reinforcing the feasibility of species-selective inhibition. The Catalophore^TM^ comparative analysis further distinguishes the binding site of Rv0183 from its human ortholog, mitigating potential cross-reactivity concerns and reinforcing the specificity of these inhibitors. Finally, we could also show that these compounds were active against live *M. tuberculosis* when grown with monoolein as the sole carbon source. These findings facilitate the next steps toward novel combination therapies aimed at treating both active and dormant tuberculosis infections.

## Acknowledgements

Parts of the work leading to these results have received support from the Innovative Medicines Initiative Joint Undertaking under Grant Agreement n′ 115489 “the European Lead Factory” and Grant Agreement No 806948 “ESCulab”, resources of which are composed of financial contribution from the European Union’s Seventh Framework Programme (FP7 /2007-2013), from the European Union’s Horizon 2020 research and innovation programme, EFPIA companies’ in kind contribution and Medicines for Malaria Venture.

This research was funded by Austrian Science Fund (FWF), Grant SFB Lipid Hydrolysis FWF F73 and by the Austrian Science Fund ÖAW APART-MINT 12033. This work was further funded by FWF, grants W901-B12, DOC46, DOC50 and DOC130, for the PhD training programs: Doctoral School - DK Molecular Enzymology, doc.fund CATalytic mechanisms and AppLications of Oxidoreductases – CATALOX, doc.fund Molecular Metabolism – MOBILES, and doc.fund Biomolecular Structures and Interactions - BioMolStruct.

We acknowledge DESY (Hamburg, Germany), a member of the Helmholtz Association HGF, for the provision of experimental facilities. Parts of this research were carried out at PETRA III and we would like to thank Johanna Hakanpää for assistance in using P11. Beamtime was allocated for proposal BAG- 20200791 EC.

The results were produced in part using high-performance computing resources and software from Innophore. Catalophore^TM^ is a registered trademark (AT 295631) of Innophore GmbH.

## Author contribution

**L.Riegler-Berket:** Conceptualization, Data curation, Formal analysis, Investigation, Methodology Project administration, Validation, Visualization, Writing – original draft, Writing – review & editing, Funding acquisition; **L.Gödl:** Conceptualization, Investigation, Methodology, Validation, Writing – original draft, Writing – review & editing; **N.Polidori:** Validation; **P.Aschauer:** Investigation, Methodology; **C.Grininger:** Investigation, Writing – review & editing; **G.Prosser:** Investigation, Methodology, Formal analysis, Writing – original draft; **J.Lichtenegger:** Investigation; **T. Sagmeister**: Validation; **L. Parigger:** Investigation, Methodology, Visualization, Writing – original draft; **C. Gruber:** Resources; **N. Reiling:** Methodology, Writing – original draft; **M. Oberer**: Conceptualization, Data curation, Formal analysis, Funding acquisition, Methodology, Project administration, Resources, Supervision, Validation, Visualization, Writing – original draft, Writing – review & editing.

## References

1. World Health Organization. *WHO - Global Tuberculosis Report* 2024. World Health Organization https://www.who.int/teams/global-tuberculosis-programme/tb-reports/global-tuberculosis-report-2024 (2024).

2. World Health Organization. *WHO - Bacterial Priority Pathogens List 2024*. https://iris.who.int/bitstream/handle/10665/376776/9789240093461-eng.pdf?sequence=1 (2024).

3. World Health Organization. Implementing the End TB Strategy: The Essentials 2022 Update. vol. 30 (2022).

4. Abrahams, K. A. & Besra, G. S. Mycobacterial drug discovery. RSC Med Chem 11, 1354–1365 (2020).

5. Dartois, V. A. & Rubin, E. J. Anti-tuberculosis treatment strategies and drug development: challenges and priorities. Nature Reviews Microbiology vol. 20 685–701 Preprint at 10.1038/s41579-022-00731-y (2022).

6. Verma, A., Ghoshal, A., Dwivedi, V. P. & Bhaskar, A. Tuberculosis: The success tale of less explored dormant Mycobacterium tuberculosis. Front Cell Infect Microbiol 12, (2022).

7. Hu, Y., Coates, A. R. M. & Mitchison, D. A. Sterilizing activities of fluoroquinolones against rifampin-tolerant populations of Mycobacterium tuberculosis. Antimicrob Agents Chemother 47, 653–657 (2003).

8. Rameshwaram, N. R., Singh, P., Ghosh, S. & Mukhopadhyay, S. Lipid metabolism and intracellular bacterial virulence: key to next-generation therapeutics. Future Microbiol 13, 1301–1328 (2018).

9. Cevik, M. et al. Bedaquiline-pretomanid-moxifloxacin-pyrazinamide for drug-sensitive and drug-resistant pulmonary tuberculosis treatment: a phase 2c, open-label, multicentre, partially randomised controlled trial. Lancet Infect Dis 24, 1003–1014 (2024).

10. Capela, R. et al. Target Identification in Anti-Tuberculosis Drug Discovery. Int J Mol Sci 24, (2023).

11. Brust, B. et al. Mycobacterium tuberculosis Lipolytic Enzymes as Potential Biomarkers for the Diagnosis of Active Tuberculosis. PLoS One 6, e25078–e25078 (2011).

12. Peyron, P. et al. Foamy Macrophages from Tuberculous Patients’ Granulomas Constitute a Nutrient-Rich Reservoir for M. tuberculosis Persistence. PLoS Pathog 4, e1000204 (2008).

13. Russell, D. G., Cardona, P. J., Kim, M. J., Allain, S. & Altare, F. Foamy macrophages and the progression of the human tuberculosis granuloma. Nat Immunol 10, 943–948 (2009).

14. Daniel, J., Maamar, H., Deb, C., Sirakova, T. D. & Kolattukudy, P. E. Mycobacterium tuberculosis uses host triacylglycerol to accumulate lipid droplets and acquires a dormancy-like phenotype in lipid-loaded macrophages. PLoS Pathog 7, (2011).

15. Babin, B. M. et al. Identification of covalent inhibitors that disrupt M. tuberculosis growth by targeting multiple serine hydrolases involved in lipid metabolism. Cell Chem Biol 29, 897–909.e7 (2022).

16. Simon, G. M. & Cravatt, B. F. Activity-based proteomics of enzyme superfamilies: Serine hydrolases as a case study. Journal of Biological Chemistry 285, 11051–11055 (2010).

17. Ortega, C. et al. Systematic Survey of Serine Hydrolase Activity in Mycobacterium tuberculosis Defines Changes Associated with Persistence. Cell Chem Biol 23, 290–298 (2016).

18. Tallman, K. R., Levine, S. R. & Beatty, K. E. Small-Molecule Probes Reveal Esterases with Persistent Activity in Dormant and Reactivating Mycobacterium tuberculosis. ACS Infect Dis 2, 936–944 (2016).

19. Li, M. et al. Identification of cell wall synthesis inhibitors active against Mycobacterium tuberculosis by competitive activity-based protein profiling. Cell Chem Biol 29, 883–896.e5 (2022).

20. Côtes, K. et al. Lipolytic enzymes in Mycobacterium tuberculosis. Appl Microbiol Biotechnol 78, 741–749 (2008).

21. Mattow, J. et al. Comparative proteome analysis of culture supernatant proteins from virulent Mycobacterium tuberculosis H37Rv and attenuated M. bovis BCG Copenhagen. Electrophoresis 24, 3405–3420 (2003).

22. Dhouib, R., Laval, F., Carriere, F., Daffe, M. & Canaan, S. A Monoacylglycerol Lipase from Mycobacterium smegmatis Involved in Bacterial Cell Interaction. J Bacteriol 192, 4776–4785 (2010).

23. Dedieu, L., Serveau-Avesque, C., Kremer, L. & Canaan, S. Mycobacterial lipolytic enzymes: a gold mine for tuberculosis research. Biochimie 95, 66–73 (2013).

24. Sassetti, C. M., Boyd, D. H. & Rubin, E. J. Genes required for mycobacterial growth defined by high density mutagenesis. Mol Microbiol 48, 77–84 (2003).

25. Bosch, B. et al. Genome-wide gene expression tuning reveals diverse vulnerabilities of M. tuberculosis Correspondence In brief ll Genome-wide gene expression tuning reveals diverse vulnerabilities of M. tuberculosis. Cell 184, 4579–4592.e24 (2021).

26. Nisa, A. et al. Different modalities of host cell death and their impact on Mycobacterium tuberculosis infection. Am J Physiol Cell Physiol 323, C1444–C1474 (2022).

27. Xu, G. et al. Hemolytic phospholipase Rv0183 of Mycobacterium tuberculosis induces inflammatory response and apoptosis in alveolar macrophage RAW264.7 cells. Can J Microbiol 56, 916–924 (2010).

28. Aschauer, P., Zimmermann, R., Breinbauer, R., Pavkov-Keller, T. & Oberer, M. The crystal structure of monoacylglycerol lipase from M. tuberculosis reveals the basis for specific inhibition. Sci Rep 8, 8948 (2018).

29. Grininger, C. et al. Structural Changes in the Cap of Rv0183/mtbMGL Modulate the Shape of the Binding Pocket. Biomolecules 11, 1299 (2021).

30. Riegler-Berket, L., Leitmeier, A., Aschauer, P., Dreveny, I. & Oberer, M. Identification of lipases with activity towards monoacylglycerol by criterion of conserved cap architectures. Biochim Biophys Acta Mol Cell Biol Lipids 1863, 679–687 (2018).

31. van Vlijmen, H. et al. The European Lead Factory: Results from a decade of collaborative, public– private, drug discovery programs. Drug Discov Today 29, 103886 (2024).

32. Schalk-Hihi, C. et al. Crystal structure of a soluble form of human monoglyceride lipase in complex with an inhibitor at 1.35 A ° resolution. Protein Science 20, 670–683 (2011).

33. Rengachari, S. et al. Conformational Plasticity and Ligand Binding of Bacterial Monoacylglycerol Lipase. Journal of Biological Chemistry 288, 31093–31104 (2013).

34. Burkhardt, A. et al. Status of the crystallography beamlines at PETRA III. The European Physical Journal Plus 131, 56 (2016).

35. Kabsch, W. Integration, scaling, space-group assignment and post-refinement. Acta Crystallogr D Biol Crystallogr 66, 133–144 (2010).

36. Krissinel, E., Uski, V., Lebedev, A., Winn, M. & Ballard, C. Distributed computing for macromolecular crystallography. Acta Crystallogr D Struct Biol 74, 143–151 (2018).

37. Winn, M. D. et al. Overview of the CCP4 suite and current developments. Acta Crystallogr D Biol Crystallogr 67, 235–242 (2011).

38. McCoy, A. J. et al. Phaser crystallographic software. J Appl Crystallogr 40, 658–674 (2007).

39. Emsley, P., Lohkamp, B., Scott, W. G. & Cowtan, K. Features and development of Coot. Acta Crystallogr D Biol Crystallogr 66, 486–501 (2010).

40. Liebschner, D. et al. Macromolecular structure determination using X-rays, neutrons and electrons: recent developments in Phenix. Acta Crystallogr D Struct Biol 75, 861–877 (2019).

41. Altschul, S. F., Gish, W., Miller, W., Myers, E. W. & Lipman, D. J. Basic local alignment search tool. J Mol Biol 215, 403–410 (1990).

42. van Kempen, M. et al. Fast and accurate protein structure search with Foldseek. Nat Biotechnol 42, 243–246 (2024).

43. Gruber, K., Steinkellner, G. & Gruber, C. Determining novel enzymatic functionalities using three-dimensional point clouds representing physico-chemical properties of protein cavities. US20150302142A1 (2015).

44. Steinkellner, G. et al. Identification of promiscuous ene-reductase activity by mining structural databases using active site constellations. Nat Commun 5, 4150 (2014).

45. Hendlich, M., Rippmann, F. & Barnickel, G. LIGSITE: automatic and efficient detection of potential small molecule-binding sites in proteins. J Mol Graph Model 15, 359–363 (1997).

46. Virtanen, P. et al. SciPy 1.0: fundamental algorithms for scientific computing in Python. Nat Methods 17, 261–272 (2020).

47. Trott, O. & Olson, A. J. AutoDock Vina: improving the speed and accuracy of docking with a new scoring function, efficient optimization, and multithreading. J Comput Chem 31, 455–461 (2010).

48. Krieger, E. & Vriend, G. New ways to boost molecular dynamics simulations. J Comput Chem 36, 996–1007 (2015).

49. Duan, Y. et al. A point-charge force field for molecular mechanics simulations of proteins based on condensed-phase quantum mechanical calculations. J Comput Chem 24, 1999–2012 (2003).

50. Hunter, J. D. Matplotlib: A 2D graphics environment. Comput Sci Eng 9, 90–95 (2007).

51. Waskom, M. L. seaborn: statistical data visualization. J Open Source Softw 6, 3021 (2021).

52. Schalk-Hihi, C. et al. Crystal structure of a soluble form of human monoglyceride lipase in complex with an inhibitor at 1.35 A ° resolution. Protein Science 20, 670–683 (2011).

53. He, Y. et al. Development of High Brain-Penetrant and Reversible Monoacylglycerol Lipase PET Tracers for Neuroimaging. J Med Chem 65, 2191–2207 (2022).

54. Aida, J. et al. Design, Synthesis, and Evaluation of Piperazinyl Pyrrolidin-2-ones as a Novel Series of Reversible Monoacylglycerol Lipase Inhibitors. J Med Chem 61, 9205–9217 (2018).

55. Labar, G. et al. Crystal structure of the human monoacylglycerol lipase, a key actor in endocannabinoid signaling. ChemBioChem 11, 218–227 (2010).

56. Bertrand, T. et al. Structural Basis for Human Monoglyceride Lipase Inhibition. J Mol Biol 396, 663–673 (2010).

57. Jiang, M. et al. A monoacylglycerol lipase inhibitor showing therapeutic efficacy in mice without central side effects or dependence. Nat Commun 14, 8039 (2023).

58. McAllister, L. A. et al. Discovery of Trifluoromethyl Glycol Carbamates as Potent and Selective Covalent Monoacylglycerol Lipase (MAGL) Inhibitors for Treatment of Neuroinflammation. J Med Chem 61, 3008–3026 (2018).

59. Butler, C. R. et al. Azetidine and Piperidine Carbamates as Efficient, Covalent Inhibitors of Monoacylglycerol Lipase. J Med Chem 60, 9860–9873 (2017).

60. Ikeda, S. et al. Design and Synthesis of Novel Spiro Derivatives as Potent and Reversible Monoacylglycerol Lipase (MAGL) Inhibitors: Bioisosteric Transformation from 3-Oxo-3,4- dihydro-2 H-benzo[b][1,4]oxazin-6-yl Moiety. J Med Chem 64, 11014–11044 (2021).

61. Jiang, M. et al. A monoacylglycerol lipase inhibitor showing therapeutic efficacy in mice without central side effects or dependence. Nat Commun 14, (2023).

62. Nawrotek, A., Talagas, A., Vuillard, L. & Miallau, L. Crystal structure of human Monoacylglycerol Lipase ABHD6 in complex with oleic acid and octyl glucoside. Worldwide Protein Data Bank (2021) doi:10.2210/PDB7OTS/PDB.

63. Pusch, L. M., Riegler-Berket, L., Oberer, M., Zimmermann, R. & Taschler, U. α/β-Hydrolase Domain-Containing 6 (ABHD6)- A Multifunctional Lipid Hydrolase. Metabolites 12, 761 (2022).

64. Dejesus, M. A. et al. Comprehensive Essentiality Analysis of the Mycobacterium tuberculosis Genome via Saturating Transposon Mutagenesis. mBio 8, e02133–16 (2017).

65. Aschauer, P. et al. Crystal structure of the Saccharomyces cerevisiae monoglyceride lipase Yju3p. Biochim Biophys Acta Mol Cell Biol Lipids 1861, 462–470 (2016).

66. Rengachari, S. et al. The structure of monoacylglycerol lipase from Bacillus sp. H257 reveals unexpected conservation of the cap architecture between bacterial and human enzymes. Biochim Biophys Acta Mol Cell Biol Lipids 1821, 1012–1021 (2012).

67. Labar, G. et al. Crystal structure of the human monoacylglycerol lipase, a key actor in endocannabinoid signaling. ChemBioChem 11, 218–227 (2010).

68. Xing, L. et al. Fluorine in Drug Design: A Case Study with Fluoroanisoles. ChemMedChem 10, 715–726 (2015).

69. Long, J. Z. et al. Selective blockade of 2-arachidonoylglycerol hydrolysis produces cannabinoid behavioral effects. Nat Chem Biol 5, 37–44 (2009).

70. Nomura, D. K. et al. Monoacylglycerol lipase exerts dual control over endocannabinoid and fatty acid pathways to support prostate cancer. Chem Biol 18, 846–856 (2011).

71. Bononi, G. et al. Reversible Monoacylglycerol Lipase Inhibitors: Discovery of a New Class of Benzylpiperidine Derivatives. J Med Chem 65, 7118–7140 (2022).

72. Muccioli, G. G., Labar, G. & Lambert, D. M. CAY10499, a novel monoglyceride lipase inhibitor evidenced by an expeditious MGL assay. Chembiochem 9, 2704–2710 (2008).

73. Granchi, C. et al. Design, synthesis and biological evaluation of second-generation benzoylpiperidine derivatives as reversible monoacylglycerol lipase (MAGL) inhibitors. Eur J Med Chem 209, 112857 (2021).

74. Jiang, M. et al. Structure-Activity Relationship Studies of Aryl Sulfoxides as Reversible Monoacylglycerol Lipase Inhibitors. J Med Chem 67, 12331 (2024).

75. Bononi, G., Lonzi, C., Tuccinardi, T., Minutolo, F. & Granchi, C. The Benzoylpiperidine Fragment as a Privileged Structure in Medicinal Chemistry: A Comprehensive Review. Molecules 29, 1930 (2024).

76. Taschler, U. et al. Monoglyceride Lipase Deficiency in Mice Impairs Lipolysis and Attenuates Diet-induced Insulin Resistance. Journal of Biological Chemistry 286, 17467–17477 (2011).

77. Steinkellner, G. et al. Identification of promiscuous ene-reductase activity by mining structural databases using active site constellations. Nat Commun 5, (2014).

78. Hetmann, M. et al. Identification and validation of fusidic acid and flufenamic acid as inhibitors of SARS-CoV-2 replication using DrugSolver CavitomiX. Sci Rep 13, (2023).

79. Shan Chang, D. P. & Guan, X. L. Metabolic Versatility of Mycobacterium tuberculosis during Infection and Dormancy. Metabolites 11, 1–21 (2021).

80. Santucci, P. et al. Delineating the physiological roles of the PE and catalytic domains of LipY in lipid consumption in mycobacterium-infected foamy macrophages. Infect Immun 86, e00394–18 (2018).

81. Santucci, P. et al. Nitrogen deprivation induces triacylglycerol accumulation, drug tolerance and hypervirulence in mycobacteria. Sci Rep 9, 8667 (2019).

82. Saxena, A. K. et al. Identification and characterisation of small-molecule inhibitors of Rv3097c- encoded lipase (LipY) of Mycobacterium tuberculosis that selectively inhibit growth of bacilli in hypoxia. Int J Antimicrob Agents 42, 27–35 (2013).

83. Deb, C. et al. A novel lipase belonging to the hormone-sensitive lipase family induced under starvation to utilize stored triacylglycerol in Mycobacterium tuberculosis. Journal of Biological Chemistry 281, 3866–3875 (2006).

