## Supplementary figures and images for "*M. tuberculosis* meets European Lead Factory – identification and structural characterization of novel Rv0183 inhibitors using X-ray crystallography"

### Supplemental Material S1

Resynthesis:

#### ELF1

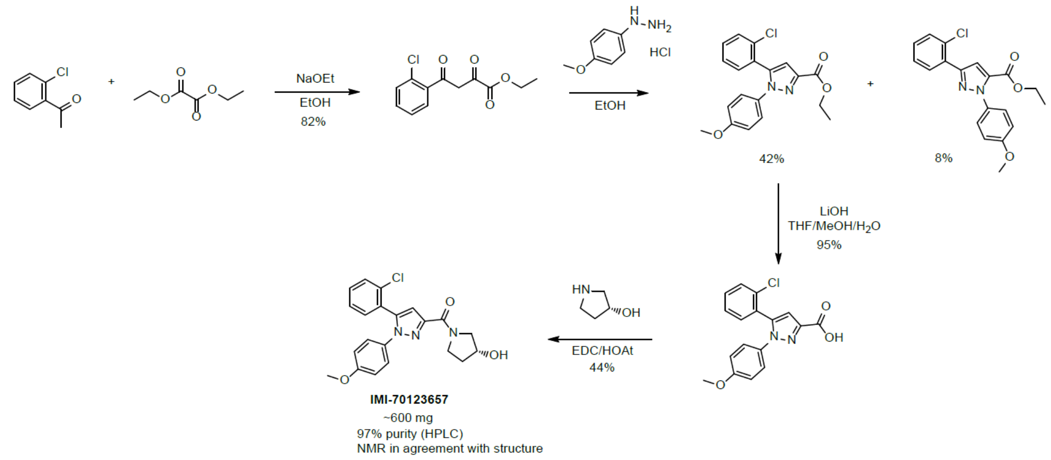


#### ELF5

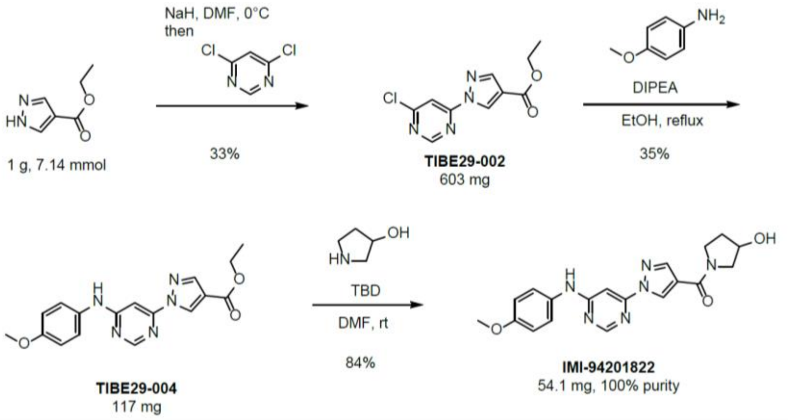


#### ELF8

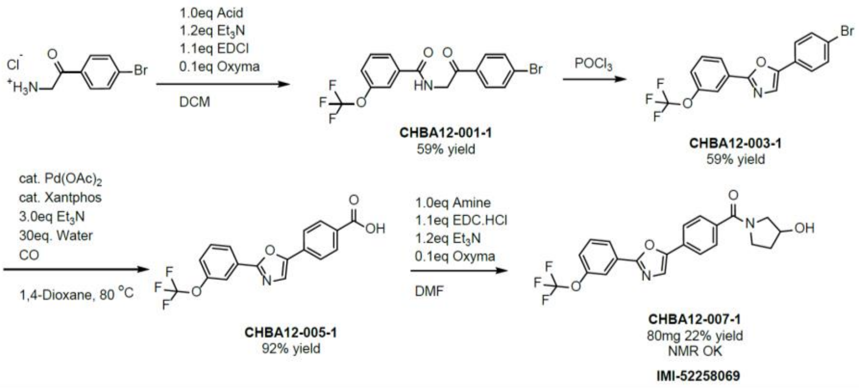
