## Supplemental Material S2 for "*M. tuberculosis* meets European Lead Factory – identification and structural characterization of novel Rv0183 inhibitors using X-ray crystallography"

EFL1

| | |
|---|---|
| title | **lebe63-006-4** |
| Method | AN_BASE.M |
| Date acquired | 29-Sep-20, 18:13:10 |
| FileName | Analysis\LCMS23_20200929_lebe63-006-4_4478.D |
| Acq. method | AN_BASE |
| Column | XSelect CSH C18 (50x2.1mm 3.5µ) |
| Flow | 0.8 ml/min, Column temp: 25°C |
| Eluent A | 10mM Ammoniumbicarbonate in water (pH 9.5) |
| Eluent B | Acetonitrile |
| Lin. gradient | t=0 min 5% B, t=4.5 min 98% B, t=6 min 98% B |
| Postrun | 2 min |
| Detection | DAD (210, 220 and 220-320nm) |
| Detection | PDA (210-320nm) |
| Detection | MSD (ESI pos/neg) mass range 100-1000 |

DAD1A, Sig=270,100 Ref=off Chromatogram

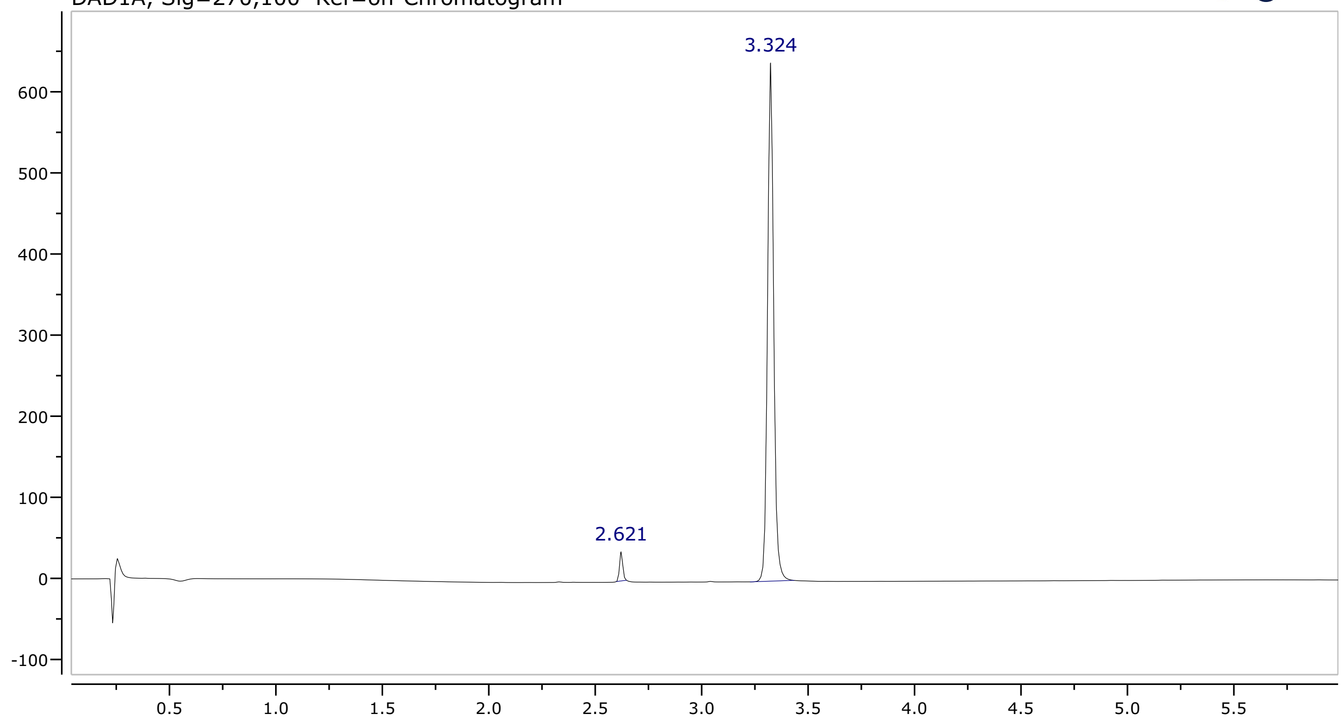


Integrals spectrum Chromatogram DAD1A, Sig=270,100 Ref=off

| rt (min) | height | area | area (%) |
|---|---|---|---|
| 2.62 | 35.98 | 752.3 | 2.84 |
| 3.32 | 639.1 | 25781 | 97.16 |

MS + spectrum 3.36

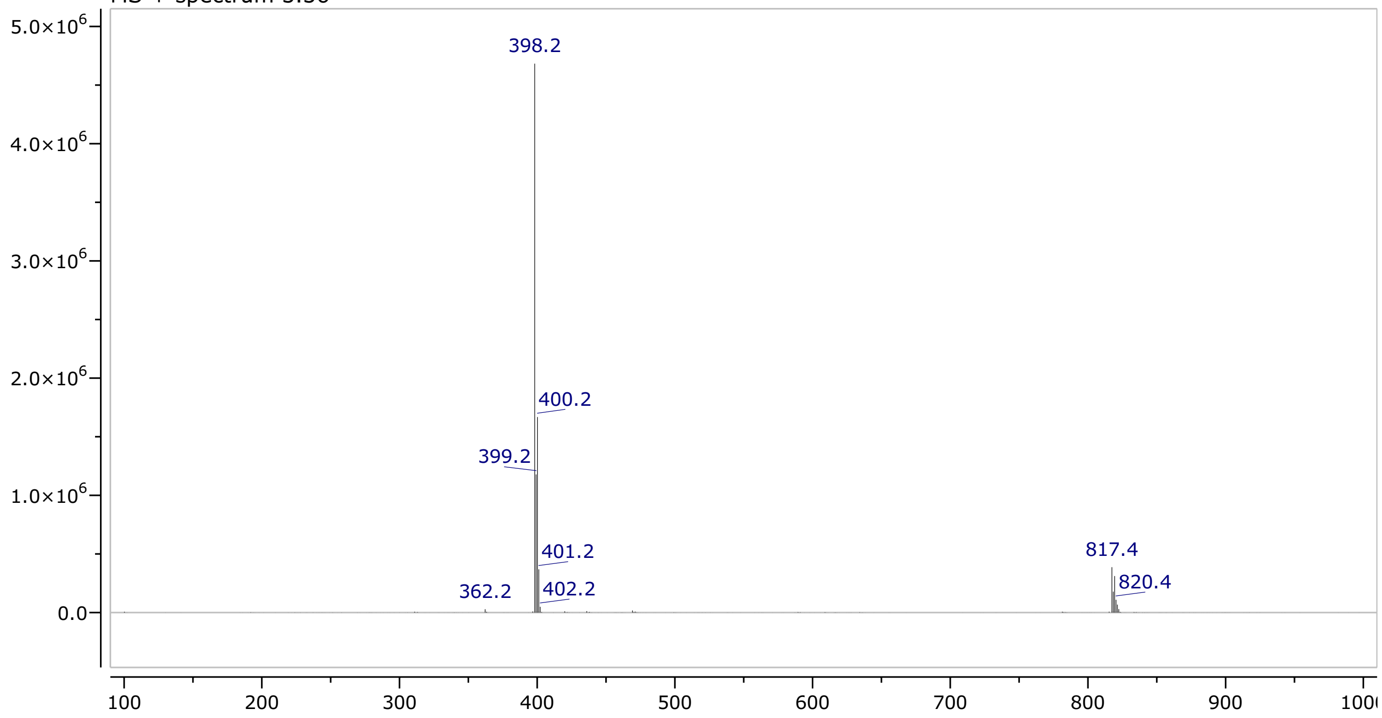

### ELF1

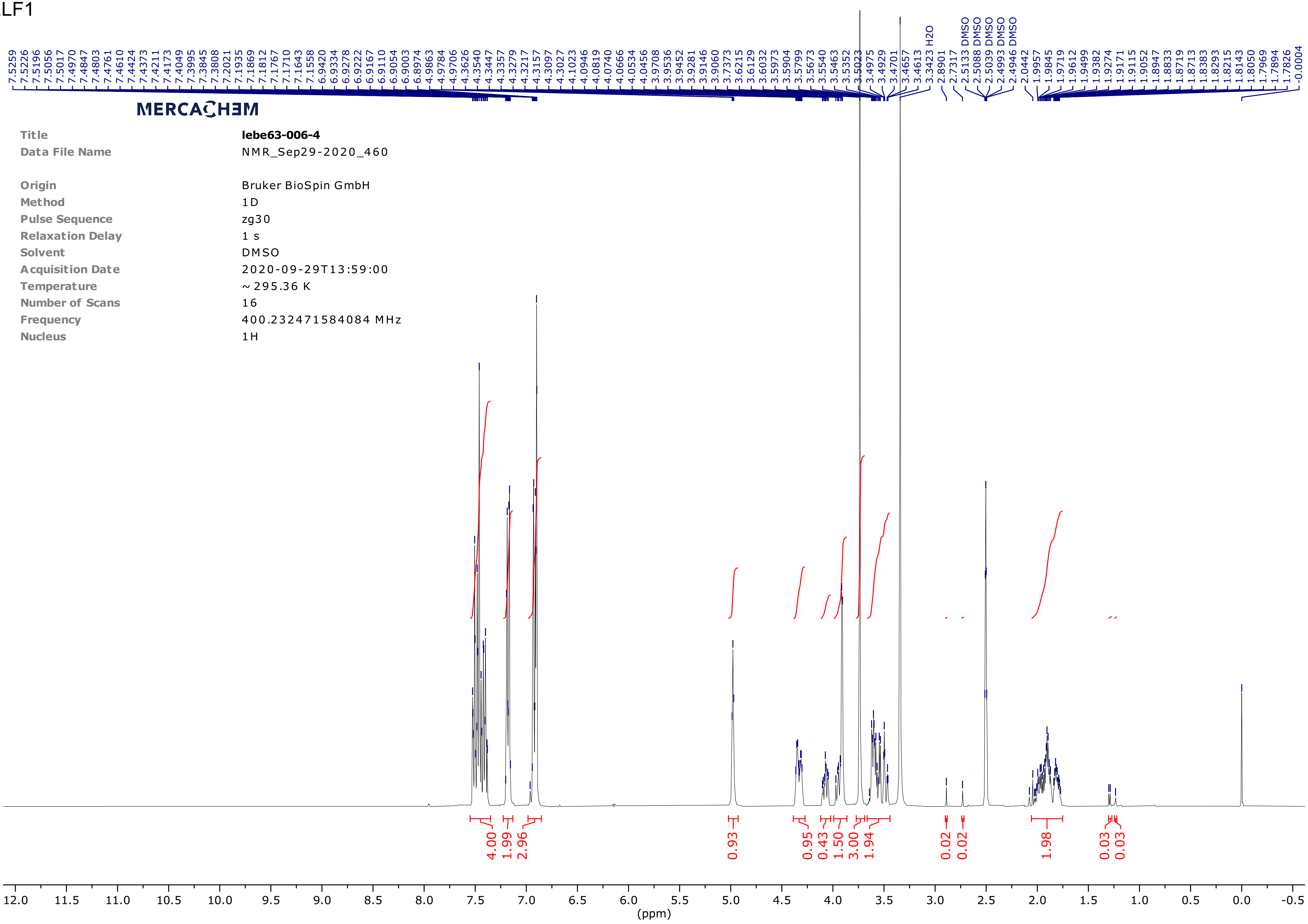


| | |
|---|---|
| Title | **lebe63-006-4** |
| Data File Name | NMR_Sep29-2020_460 |
| Origin | Bruker BioSpin GmbH |
| Method | 1D |
| Pulse Sequence | zg30 |
| Relaxation Delay | 1 s |
| Solvent | DMSO |
| Acquisition Date | 2020-09-29T13:59:00 |
| Temperature | ~ 295.36 K |
| Number of Scans | 16 |
| Frequency | 400.232471584084 MHz |
| Nucleus | 1H |

ELF5

| | |
|---|---|
| title | **TIBE29-009-1** |
| Date acquired | 08-Oct-2020, 08:25:41 |
| FileName | Analysis\LCMS20_201008_TIBE_ESCL_final_003.raw |
| UPLC | Waters I-Class |
| Acq. Method | UPLC_AN_BASE |
| Column | XSelect CSH C18 (50x2.1mm 2.5µm) |
| Eluent A | 10mM ammoniumbicarbonate in water (pH 9.5) |
| Eluent B | acetonitrile |
| Gradient | t=0 min 5% B, t= 2.0 min 98% B, t=2.7 min 98% B |
| Posttime | 0.3 min |
| Detection PDA | 210-320nm |
| MS | QDa |
| Detection MS | ESI (scan) |
| Mass range | 100-800 (pos) |
| Mass range | 100-800 (neg) |
| Time | 2.7 min |
| Cone | 15 V; Freq: 2 Hz |

MERCACHEM

PDA - Total Absorbance Chromatogram

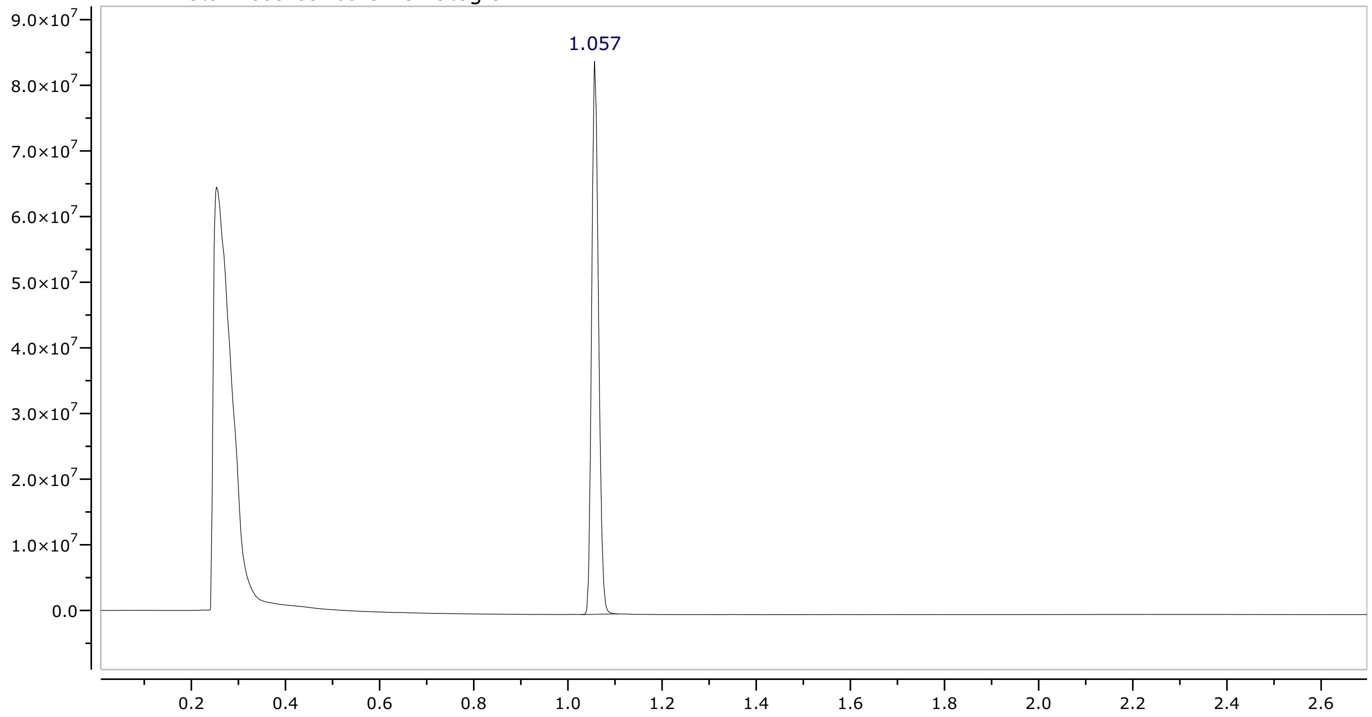


Integrals spectrum PDA - Total Absorbance Chromatogram

| rt (min) | height | area | area (%) |
|---|---|---|---|
| 1.06 | 84253851 | 1717910278 | 100.00 |

MS + spectrum 1.06

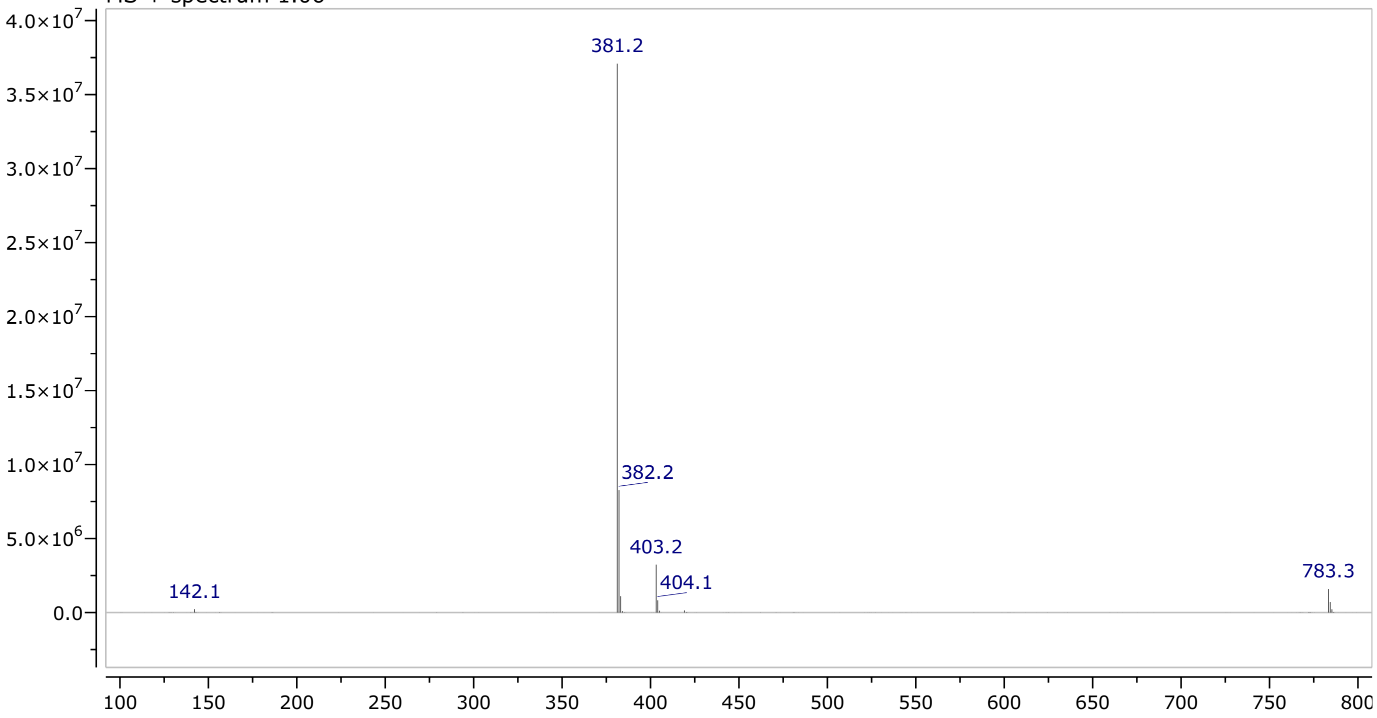

ELF5

MERCACHEM

| | |
|---|---|
| Title | **TIBE29-009-1** |
| Data File Name | NMR_Oct09-2020_490 |
| Origin | Bruker BioSpin GmbH |
| Method | 1D |
| Pulse Sequence | zg30 |
| Relaxation Delay | 1 s |
| Solvent | DMSO |
| Acquisition Date | 2020-10-09T13:13:00 |
| Temperature | ~ 294.86 K |
| Number of Scans | 16 |
| Frequency | 400.232471584084 MHz |
| Nucleus | 1H |

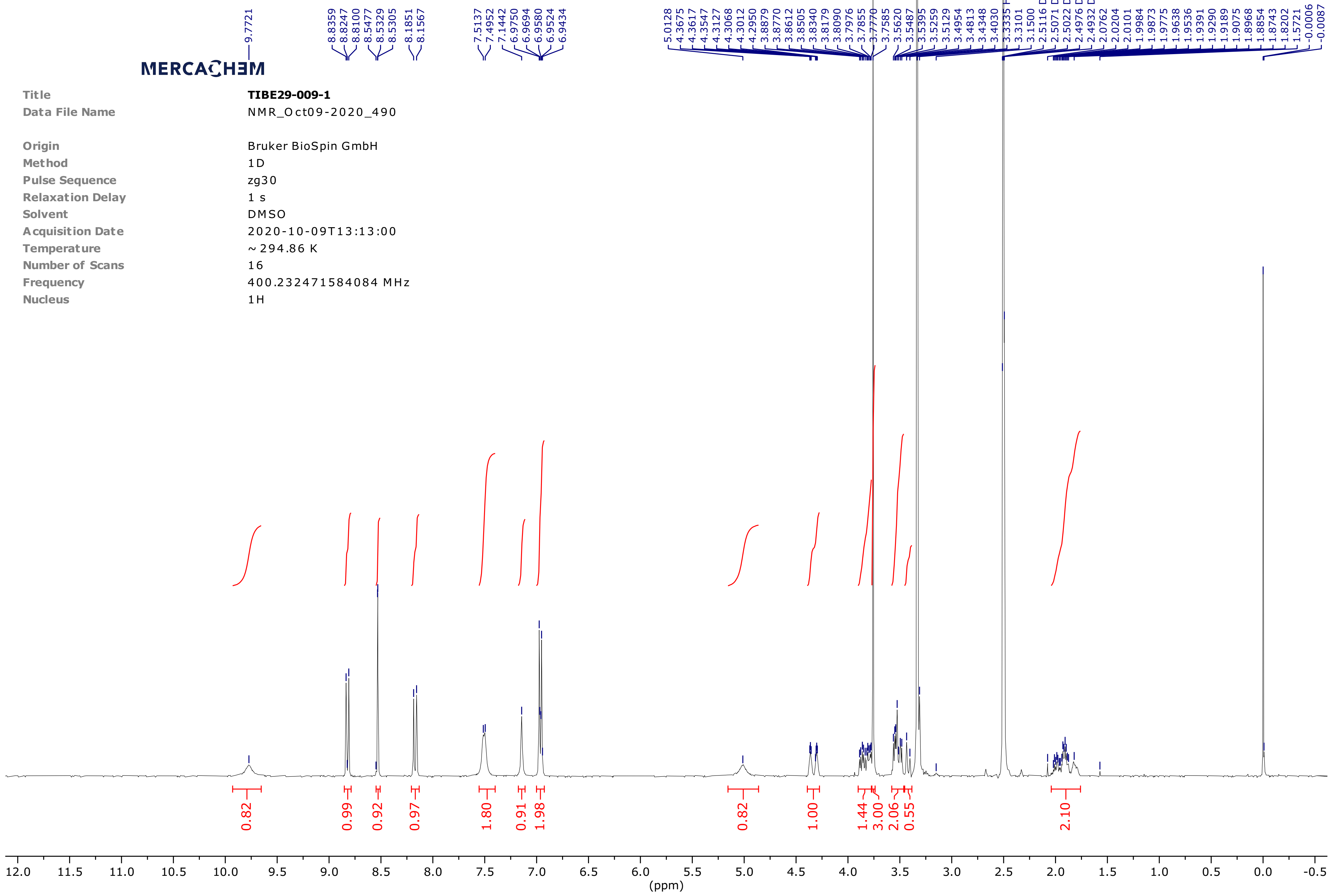

ELF8

| | |
|---|---|
| title | **CHBA12-007-1** |
| Method | AN_BASE.M |
| Date acquired | 19-Oct-20, 18:58:29 |
| FileName | Analysis\LCMS23_20201019_CHBA12-007-1_5765.D |
| Acq. method | AN_BASE |
| Column | XSelect CSH C18 (50x2.1mm 3.5μ) |
| Flow | 0.8 ml/min, Column temp: 25°C |
| Eluent A | 10mM Ammoniumbicarbonate in water (pH 9.5) |
| Eluent B | Acetonitrile |
| Lin. gradient | t=0 min 5% B, t=4.5 min 98% B, t=6 min 98% B |
| Postrun | 2 min |
| Detection | DAD (210, 220 and 220-320nm) |
| Detection | PDA (210-320nm) |
| Detection | MSD (ESI pos/neg) mass range 100-1000 |

MERCACHEM

DAD1A, Sig=270,100 Ref=off Chromatogram

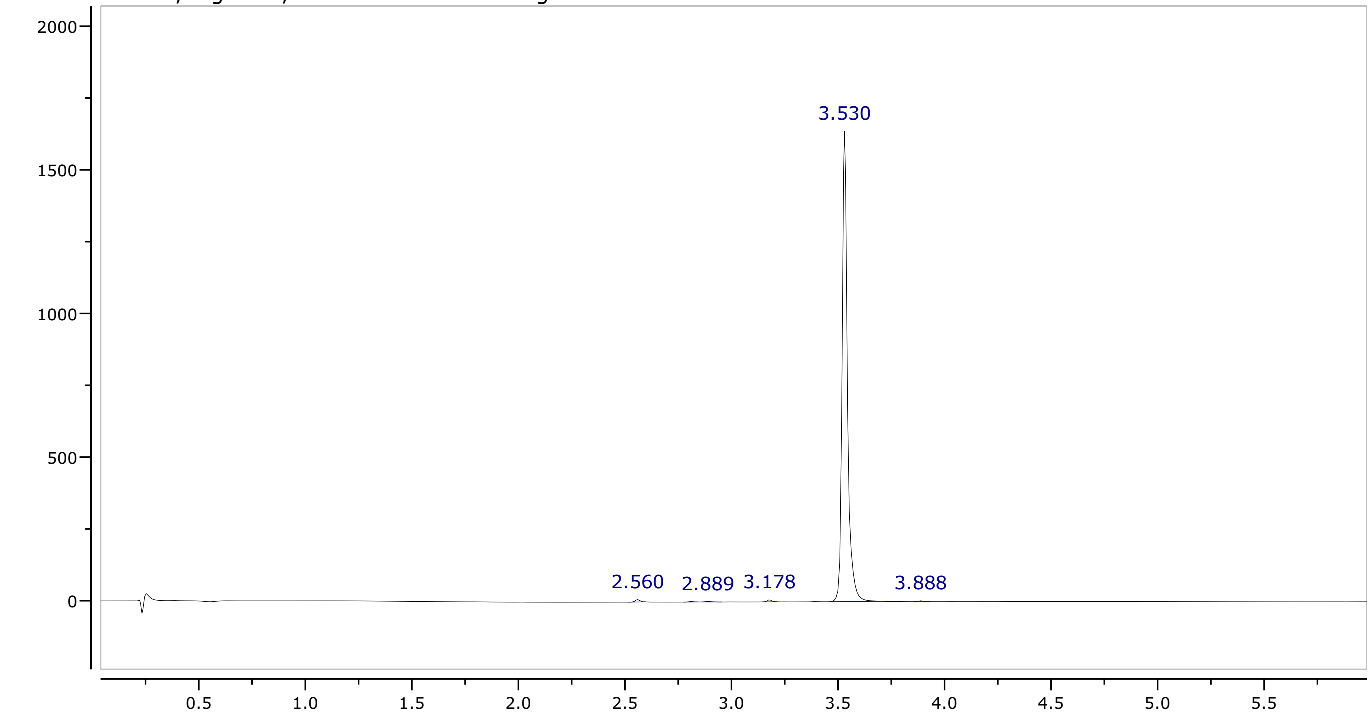


Integrals spectrum Chromatogram DAD1A, Sig=270,100 Ref=off

| rt (min) | height | area | area (%) |
|---|---|---|---|
| 2.56 | 8.459 | 251.4 | 0.43 |
| 2.81 | 1.935 | 49.76 | 0.09 |
| 2.89 | 1.967 | 73.25 | 0.13 |
| 3.18 | 7.201 | 195.6 | 0.34 |
| 3.53 | 1637 | 57412 | 98.88 |
| 3.89 | 2.749 | 82.25 | 0.14 |

MS + spectrum 3.55

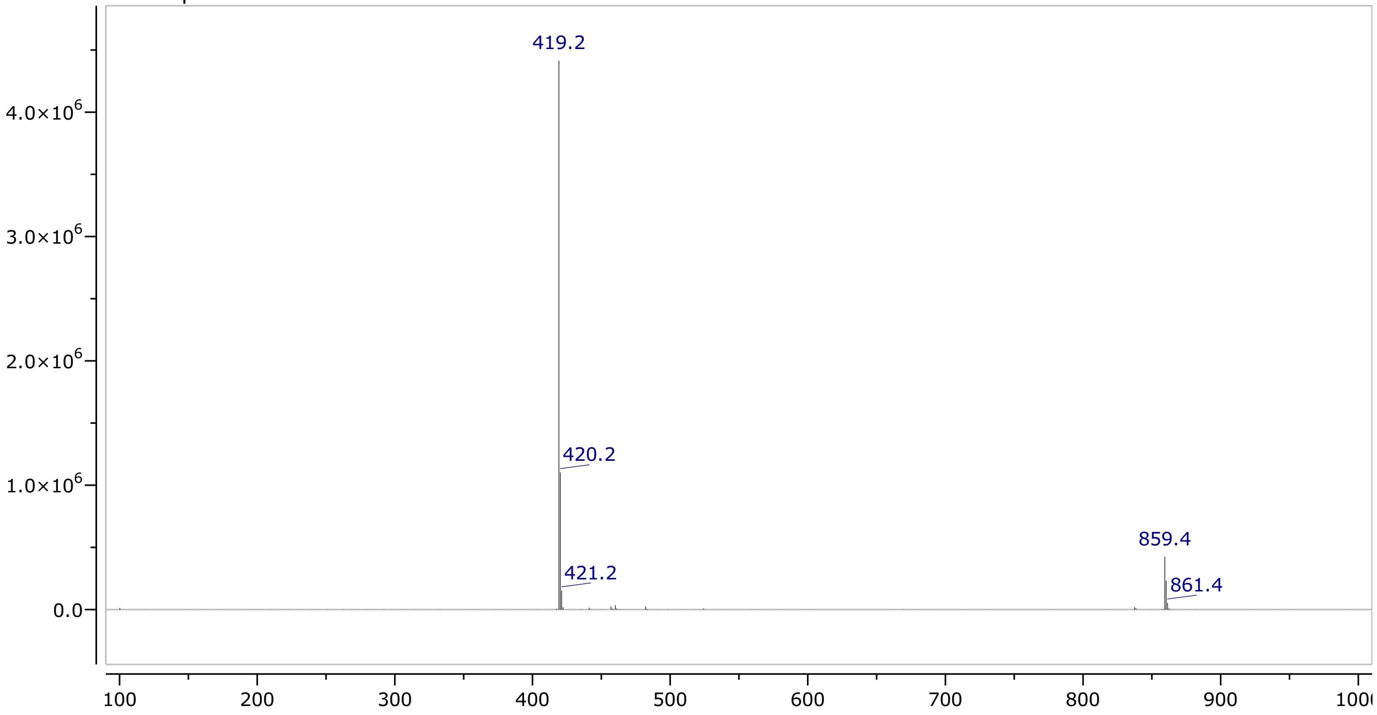

ELF8

MERCACHEM

| | |
|---|---|
| Title | **CHBA 12-007-1** |
| Data File Name | NMR-RUN_1019_135647_440 |
| Origin | Bruker BioSpin GmbH |
| Method | 1D |
| Pulse Sequence | zg30 |
| Relaxation Delay | 1 s |
| Solvent | DMSO |
| Acquisition Date | 2020-10-19T14:02:00 |
| Temperature | ~ 295.9502 K |
| Number of Scans | 64 |
| Frequency | 400.132470966543 MHz |
| Nucleus | 1H |

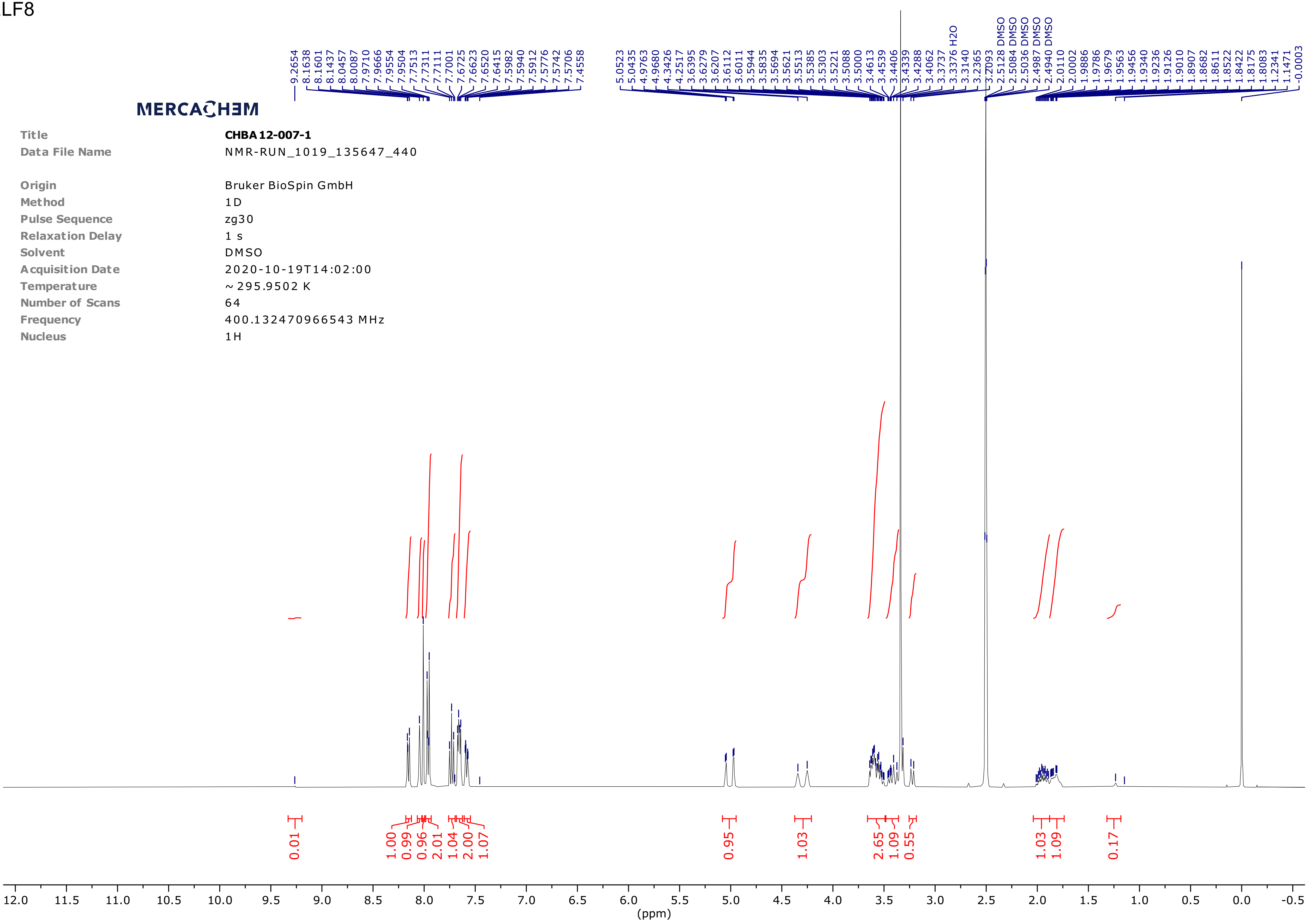
